# Cryo-EM structures of Kv1.2 potassium channels, conducting and non-conducting

**DOI:** 10.1101/2023.06.02.543446

**Authors:** Yangyu Wu, Yangyang Yan, Youshan Yang, Shumin Bian, Alberto Rivetta, Ken Allen, Fred J. Sigworth

## Abstract

We present near-atomic-resolution cryo-EM structures of the mammalian voltage-gated potassium channel Kv1.2 in open, C-type inactivated, toxin-blocked and sodium-bound states at 3.2 Å, 2.5 Å, 3.2 Å, and 2.9Å. These structures, all obtained at nominally zero membrane potential in detergent micelles, reveal distinct ion-occupancy patterns in the selectivity filter. The first two structures are very similar to those reported in the related Shaker channel and the much-studied Kv1.2-2.1 chimeric channel. On the other hand, two new structures show unexpected patterns of ion occupancy. First, the toxin α-Dendrotoxin, like Charybdotoxin, is seen to attach to the negatively-charged channel outer mouth, and a lysine residue penetrates into the selectivity filter, with the terminal amine coordinated by carbonyls, partially disrupting the outermost ion-binding site. In the remainder of the filter two densities of bound ions are observed, rather than three as observed with other toxin-blocked Kv channels. Second, a structure of Kv1.2 in Na^+^ solution does not show collapse or destabilization of the selectivity filter, but instead shows an intact selectivity filter with ion density in each binding site. We also attempted to image the C-type inactivated Kv1.2 W366F channel in Na^+^ solution, but the protein conformation was seen to be highly variable and only a low-resolution structure could be obtained. These findings present new insights into the stability of the selectivity filter and the mechanism of toxin block of this intensively studied, voltage-gated potassium channel.

## Introduction

Among the most-studied ion channel proteins is the Kv1 family of tetrameric, six-transmembrane-segment, voltage-gated potassium (Kv) channels. The best-known member of this family is the *Drosophila* Shaker channel, first cloned in the late 1980s (Iverson et al., 1988; Papazian et al., 1987) and which quickly became the favorite object for biophysical studies due to its small size and ease of heterologous expression. The potassium current in the squid giant axon described by Hodgkin and Huxley (Hodgkin & Huxley, 1952) is carried by a closely related Shaker homologue (Rosenthal & Bezanilla, 2002).

In 2005 Long et al. (2005) obtained the 2.9 Å crystal structure of Kv1.2, a mammalian Shaker-family channel. This structure showed clearly the overall architecture of the channel, but unfortunately the densities were weak in the all-important voltage-sensor domains (VSDs). Two years later Long et al. (Long et al., 2007) presented the 2.4Å X-ray structure of the “paddle chimera” Kv1.2-2.1 channel in which a portion of the voltage-sensor domain (the S3-S4 paddle) was replaced by the corresponding Kv2.1 sequence. This structure presented the VSDs in fine detail, and so was a watershed in the structural understanding of the many functional measurements of Shaker channels. It has also formed the basis of computational studies of voltage-sensing and gating mechanisms. A subsequent cryo-EM structure of Kv1.2-2.1 channels reconstituted into nanodiscs (Matthies et al., 2018) has excellent agreement with the X-ray structure. Meanwhile, a very welcome recent development is the cryo-EM structure of the original Shaker channel itself (Tan et al., 2022).

The high-resolution structure of the Kv1.2 paddle chimera channel has been the basis for much research, but the native Kv1.2 channel has been used for more functional studies (Ishida et al., 2015; Suarez-Delgado et al., 2020; Wu et al., 2022). The properties of the native and chimeric channels are similar but not identical (Tao & MacKinnon, 2008). The chimeric channel has 27 amino-acid differences in the S3-S4 paddle region, but the VSD structures of Long et al. (2005 and 2007) are remarkably similar; in the transmembrane region the !-carbon traces are nearly superimposable. Our structure of Kv1.2 confirms this remarkable structural conservation.

Inactivation is an important process that diminishes voltage-gated channel current even when the activating membrane depolarization is maintained. A relatively slow inactivation process, usually called C-type inactivation (Hoshi et al., 1991) involves conformational changes in the pore domain and the selectivity filter. Various mutations in the pore domain have been shown to accelerate or impede C-type inactivation. The Shaker pore-domain mutation W434F renders the channel almost completely non-conductive through a process like C-type inactivation (Perozo et al., 1993; Pless et al., 2013; Suarez-Delgado et al., 2020; Yang et al., 1997). Structures of this mutant (Tan et al., 2022) and of a corresponding mutant Kv1.2-2.1 paddle-chimera channel (Reddi et al., 2022) show a dramatic dilation of the selectivity filter that disrupts two of the four ion-binding sites in the selectivity filter. We report here the structure of the corresponding W366F mutant in the Kv1.2 channel background, which shows a nearly identical dilation.

An important aspect of C-type inactivation is that it is enhanced when potassium ions are absent. In KcsA the replacement of most K^+^ with Na^+^ results in a collapse of the selectivity filter region, occluding the two central ion-binding sites (Zhou et al., 2001). Other channels including NaK2K and members of the K2P family show a destabilization of the external part of the pore at low K^+^ concentrations, abolishing the two outer ion-binding sites (Lolicato et al., 2020; Matamoros et al., 2023; Sauer et al., 2011). In low potassium the NaK2K channel actually shows a dilation of the outer selectivity filter very similar to that seen in Shaker-W434F and the corresponding K1.2-2.1 mutant. On the other hand, a lengthy molecular-dynamics simulation of deactivation in the Kv1.2-2.1 chimera channel in symmetrical K^+^ solutions showed that, with the closing of the channel gate, there is a complete loss of water and ions from the cavity below the selectivity filter. Under these conditions however the selectivity filter remains intact and is populated by K^+^ ions (Jensen et al., 2012); M. Ø. Jensen, personal communication). In view of the variety of effects of low K^+^, in our current study we sought to observe experimentally the structure of the Kv1.2 channel when potassium ions are absent.

Finally, Kv channels are targets for many toxins that inhibit channel conductance, with a common mechanism being occlusion of the pore. This occlusion has been seen directly in two cases. Crystal structures of the Kv1.2 paddle-chimera channel have been obtained with charybdotoxin (CTx) bound (Banerjee et al., 2013), and cryo-EM structures of Kv1.3 with the ShK toxin bound (Selvakumar et al. 2022). In the present work we consider dendrotoxins (DTxs) produced by the mamba snake *Delodiaspis*. These are distinct from CTx but also block Kv channels with high potency and selectivity (Gasparini et al., 1998). In this study we have determined the structure of Kv1.2 with α-DTX bound.

## Results

### Overview of mammalian Kv1.2 structure

For this study we employed a full-length Kv1.2 α-subunit construct containing three mutations in disordered regions of the N-terminus and S1-S2 linker. We denote this construct used for structure determination Kv1.2_s_. We co-expressed a construct of of Kvβ2 (Gulbis et al., 1999) containing residues 36 to 367. Each β2-subunit contained five mutations chosen to neutralize positive charges on the cytoplasmic face of the subunit; these remove strong, interfering interactions with cryo-EM carbon substrates. We find that the mutations have no effect on β-subunit secondary structure. Although α and β subunits were co-expressed, in cryo-EM micrographs we observed many α_4_ complexes along with the expected α_4_β_4_ particles. As the structures of the transmembrane regions of these two populations of channels were indistinguishable, we combined the two particle sets. This is reasonable as the presence of β2 subunits has very little effect on the structure of the α-subunit T1 domains to which they bind (Gulbis et al., 2000) and also little effect on channel function (Rettig et al., 1994).

The constructs encoding the Kv1.2_s_ or Kv1.2_s_-W366F mutant α-subunits, along with the β2 subunits, were expressed in *Pichia pastoris* essentially as described (Long et al., 2005). Channel complexes were affinity-purified in the presence of dodecylmaltoside detergent and subjected to size-exclusion chromatography in buffers containing either 150 mM K^+^ or 150 mM Na^+^ ions. The channel complexes were plunge-frozen on grids for cryo-EM analysis. Focusing on the transmembrane region of the Kv1.2 channel complexes, we obtained structures of nominally open, C-type inactivated, DTx-bound, and K^+^ free states at resolutions of 3.2 Å, 2.5 Å, 3.2 Å and 2.9 Å respectively. Most main-chain and sidechain densities were clearly visible in the resulting maps allowing atomic models to be built with high confidence.

Structures of the Kv1.2_s_ channel (Figure 1A-C, and Figure 1—figure supplement 1) are very similar to those of the Kv1.2-2.1 chimera (Long et al., 2007) and *Drosophila* Shaker (Tan et al., 2022) channels. As observed in other Kv1 and Kv2 structures, the voltage-sensing domains (VSDs) contain the membrane-spanning S1-S4 helixes (Figure 1D and E) while the S5, S6, pore helix and selectivity filter P-loop form the ion-conduction pore in a domain-swapped configuration. Densities of lipids bound to the VSDs and pore domains (PDs) are clearly visible in the Kv1.2_s_ maps (Figure 1A).

**Figure 1.**
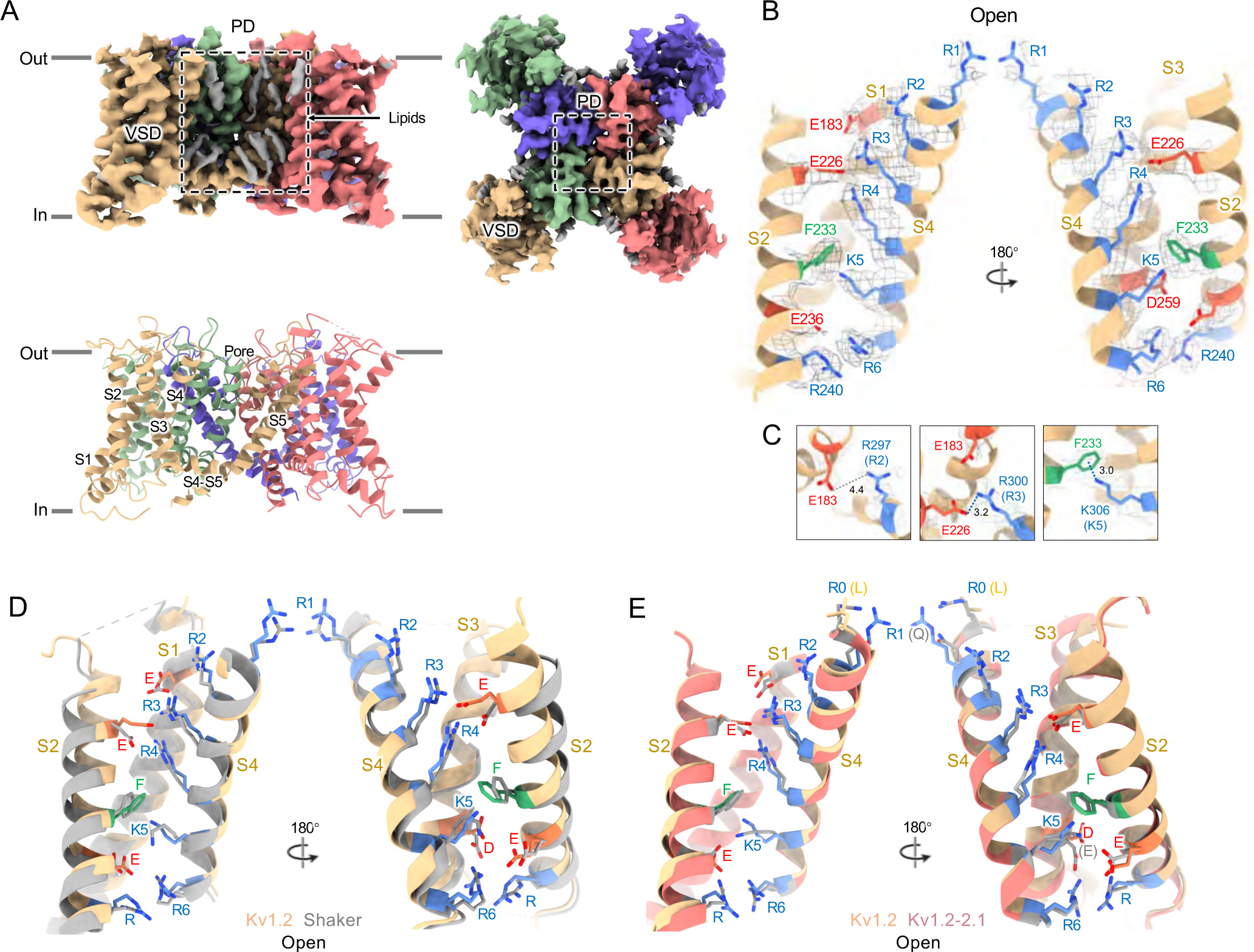
Open Kv1.2 overall structure. (A) Side and top view of the Kv1.2_s_ cryo-EM density map (upper panel) and model (lower panel). Lipid densities are colored gray in the map. (B) Side view of VSD structure with map density of Kv1.2_s_. (C) The relative positions of the interacting residues R2 and E138 (upper), R3 and E226 (middle), K5 and F233 (lower) are shown. (D) Superposition showing the very close match of Kv1.2 (yellow) and Shaker (gray) VSD structures. (E) Superposition of Kv1.2_s_ (yellow) and Kv1.2-2.1 (pink) VSD structures. Positively charged, negatively charged and aromatic residues are shown as blue, red and green, respectively. VSD, voltage-sensing domain; PD, Pore domain.

In the critical region of the voltage-sensor domain (VSD) the sidechains of the voltage-sensing Arg and the coordinating Glu and Asp residues, as well as the charge-transfer center phenylalanine (Tao et al., 2010) are essentially superimposable with Shaker (Figure 1D), with an RMS difference of all Arg sidechain atoms of 0.85 Å. As Kv1.2 is a member of the Shaker potassium channel family and has 68% amino-acid identity with *Drosophila* Shaker, it is not surprising that the fold is very similar.

As in the other homologues an open S6 gate is visible in the detergent-solubilized Kv1.2_s_ structure; this is expected for the open conformation at zero membrane potential. Seeing no evidence of a barrier to ion flux, we call this the open-channel structure (Figure 2B).

**Figure 2.**
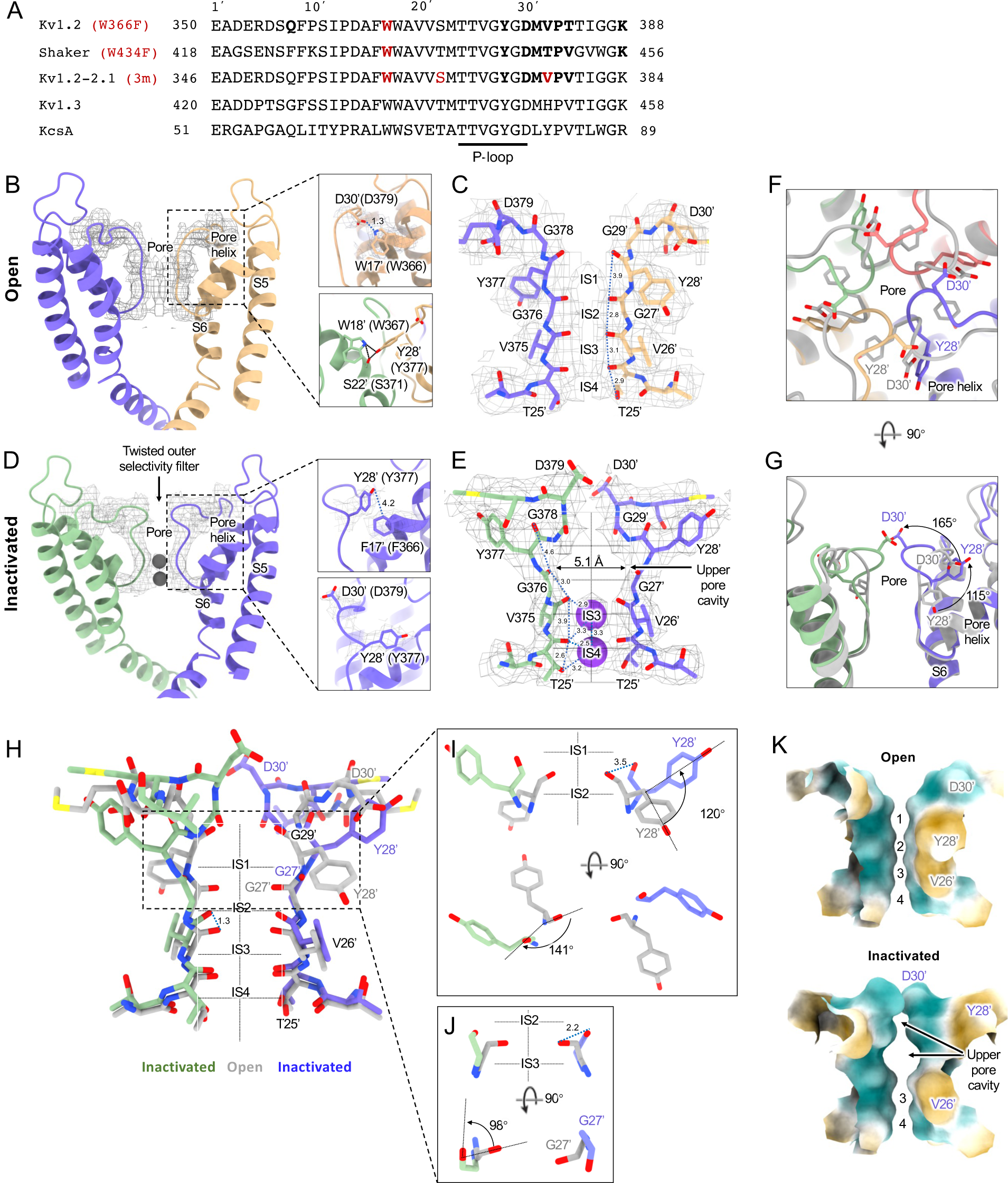
Kv1.2 pore domain and selectivity filter structures in inactivated and open conformations. (A) Sequence alignment of potassium channels in the S5-S6 linker region, with the relative numbering of Miller (1990) indicated at top. Locations of mutations of interest are noted in red. (B) Side view of opposing subunits in the open Kv1.2_s_ pore domain. Relative positions of Y28’, W17’ and D30’ are shown in the right-hand panels. Shown with a dashed blue line is the key hydrogen bond between D30’ and W17’ that is eliminated in the W17’F (W366F) mutant. (C) Side view of opposing Kv1.2_s_ P-loops. Labels on the left show the Kv1.2 residue numbering, and on the right the Miller numbering. (D) Side view of the inactivated Kv1.2_s_ W17’F pore domain. The new locations of Y28’, F17’ and D30’ are shown in the right panels. (E) Side view of the Kv1.2_s_ P-loop in the inactivated conformation. In the large upper-pore cavity the G27’ and G29’ carbonyls are 5.1 and 11 Å apart. Potassium ions are shown as purple balls. (F) Superimposed top views of the Kv1.2_s_ outer pore in inactivated (colored) and open (grey) states. In the inactivated channel the displaced Y28’ ring of one subunit is in the position occupied by D30’ - in the neighboring subunit - of the open channel. (G) The corresponding side view. The large rotation of the Y28’ side chain and flipping of D30’ are indicated by curved arrows. (H) Side view of the P-loop in inactivated (colored) and open (grey) conformations. (I,J) Details of Y28’ G27’ carbonyl and side chain reorientation from open (grey) to inactivated (colored) states are shown in side view (upper) and top view. (K) Surface renderings of open and inactivated selectivity filter region. Hydrophilic and hydrophobic surfaces are shown in teal and orange, respectively.

Despite the very closely matched open-state structures of the VSDs it is therefore puzzling that the total gating charge movement in Shaker channels is larger, about 13 elementary charges per channel, while it is only 10 e_0_ per channel in Kv1.2 (Islas, 2016). The most likely explanation is that in Kv1.2 there is some additional restraint, lacking in Shaker, on the physical displacement of each S4 helix.

In the paddle chimera a region including the N-terminal half of the S4 helix in Kv1.2 is replaced by the corresponding Kv2.1 sequence. One difference in the voltage-sensing residues is that a glutamine replaces the arginine at the first position (R1) in the chimera. Another difference is that there is an earlier Arg residue R0 that replaces leucine, but is unlikely to contribute to charge movement. A direct measurement of charge movement in the paddle chimera is lacking, but macroscopic activation and inactivation show lower voltage sensitivity (Tao & MacKinnon, 2008) as if its gating charge movement were reduced. Nevertheless, as was observed by Long et al. (2007), the structures of the S4 regions are very similar: comparing Kv1.2 with the paddle chimera (Figure 1E) the rms deviation of Arg sidechain atoms is less than 0.75 Å.

### Channel inactivation

Figure 2 compares the structure of the open channel with that of the Kv1.2_s_-W366F mutant. While the corresponding mutation W434F in Shaker essentially abolishes ion current through the channel (Yang et al., 1997) the Kv1.2_s_ mutant allows currents to flow transiently on depolarization, but decay to less than 3% of the peak within 80 ms (Figure 2—figure supplement 1). Inactivation in these channels can be slowed by raising extracellular K^+^ or by applying tetraethylammonium ion (Suarez-Delgado et al., 2020), properties that are hallmarks of C-type inactivation. Here we shall call the cryo-EM structure of Kv1.2_s_-W366F the “inactivated channel” structure, keeping in mind that other distinct inactivated channel conformations are likely to exist. The overall conformational difference between open and inactivated structures in Kv1.2s is identical to the difference observed in structures of Shaker (Tan et al., 2022) and in structures of Kv1.2-2.1 (Long et al. 2007; Reddi et al., 2022).

For detailed comparison with structures of other voltage-gated potassium channels, we borrow from Miller (1990) the numbering of the 39 residues in the S5-S6 linking region, which we will denote as residue numbers 1’ to 39’. (Figure 2A). The 39-residue numbering is appropriate for Kv1 through Kv4 channels (Shealy et al., 2003) and is fortuitously convenient in comparisons with KcsA, because it differs numerically by exactly 50 from the residue numbering of KcsA. In the Miller system for example the Kv1.2 W366 residue is W17’, and we can denote the W366F mutant channel as Kv1.2_s_-W17’F. In KcsA the W17’ residue is W67.

The selectivity filter of potassium channels consists of an array of four copies of the extended loop (the P-loop) formed by a highly conserved sequence, in this case TTVGYGD. The main-chain carbonyl oxygens of the residues TVGY, positions 25’ to 28’, delineate ion binding sites S3, S2 and S1, respectively, while the lower (most intracellular) edge of binding-site S4 is formed by the T25’ hydroxyl. Hydrogen bonds involving two P-loop residues anchor the outer half of the selectivity filter and are particularly important in inactivation mechanisms (Figure 2B, right panels) (Pless et al., 2013; Sauer et al., 2011). Normally, the tyrosine Y28’ (Y377 in Kv1.2) is constrained by hydrogen bonds to residues in the pore helix and helix S6 and is key to the conformation of the selectivity filter. The final aspartate of the P-loop, D30’ (D379 in Kv1.2) is normally located near the extracellular surface and has a side chain that also participates in H-bonds with W17’ (W366 in Kv1.2) on the pore helix.

The difference between the open and inactivated Kv1.2_s_ structures, like the open-inactivated state differences in Kv1.2-2.1 (Reddi et al., 2022) and in Shaker (Tan et al., 2022) can be imagined as resulting from a two-step process. The first step is a partial twist of the P-loop backbone involving D30’. When W17’ is mutated to phenylalanine, it is no longer an H-bond donor to D30’ (Figure 2D right upper panel). The resulting destabilization of D30’ allows it to reorient toward the external water-filled vestibule.

The second step is the reorientation of Y28’ and further twisting of the polypeptide backbone. Y28’ normally participates in H-bonds with the pore-helix residues W18’ and S22’ (Figure 2B, right-hand panels), but with the release of D30’, and presumably the entry of water molecules into the space surrounding the P-loop, the side chain of Y28’ also reorients toward the external solution, filling some of the original volume occupied by the side chain of D30’. The reorientation of the phenol group of Y28’ is through a very large pitch angle of about 120° (Figure 2G,H,I). The reorientations of the D30’ and Y28’ side chains drag and twist the backbone. The result is an enlargement of the ion-binding site IS2 to a width of 5.1Å between opposing carbonyl oxygens (Figure 2E) and an even wider (11 Å) cavity is formed at the level of IS1. There are no clear ion densities in the upper selectivity filter.

Meanwhile, the lower binding sites IS3 and IS4 show relatively small disturbances in the inactivated state. The largest change is a rise of 1.3Å in the position of the G27’ carbonyl, the one which defines the top of IS3 (Figure 2J). As in the inactivated Shaker structure (Tan et al., 2022) strong ion densities are seen in IS3 and IS4.

Might symmetry-breaking accompany Kv1.2 inactivation? MD simulations of inactivated paddle chimera, KcsA and hERG channels (Kondo et al., 2018; Li, Shen, Reddy, et al., 2021; Li, Shen, Rohaim, et al., 2021), all homotetramers, show a change to two-fold symmetry in the pore region such that the selectivity filter has distinct conformations for each pair of opposing subunits. However, we were not able to detect any evidence of reduced symmetry in the inactivated Kv1.2 channel. We were able to build the atomic model unambiguously into the 2.5 Å map that was obtained with C4 symmetry imposed. Further, there was no evidence of broken symmetry when using symmetry expansion and local C1 refinements of the cryo-EM dataset.

A comparison of our open-channel and inactivated Kv1.2_s_ structures show subtle but noticeable differences in the VSDs. Salt bridges involving the S4 Arg and Lys residues are shifted slightly (Figure 2—figure supplement 3A-D). Arg300 (R3) is in close proximity to Glu226 on the S2 helix for the open channel, while R3 is closer to Glu183 in the S2 helix. The Glu226 side chain adopts a visible interaction with R4 in the inactivated state. In both open and inactivated states of Kv1.2_s_, K5 interacts more closely with S2 Phe233 than is the case in the paddle chimera channel (Figure 1E).

Functional interactions have been observed between residues in the voltage-sensing domains and pore residues involved in C-type inactivation (Bassetto et al., 2021; Conti et al., 2016). Leucine and valine residues on the S4 helix interface with serine and leucine residues on S5, which in turn contact F16’ and W17’ on the pore helix. We see very little relative movement of the residues at the S4-S5 interface, much less than 1Å difference between open and inactivated structures.

When individual VSD domains are aligned the VSD helices in Kv1.2_s_ and the inactivated Kv1.2_s_-W17’F superimpose very well at the top (including the S4-S5 interface described above), there is a general twist of the VSD helix bundle that yields an overall rotation of about 3° at the bottom of the bundle (Figure 2—figure supplement 3E-F). In moving to the inactivated state, the axes of the helices S0, S1 and S2 tilt by 6°, 4° and 3.2° in a clockwise direction as viewed from the cytoplasmic side, while S4 is stationary. This lower VSD rotation provides a good explanation of the shifts in the S2 residues Glu183 and Glu226 that interact with R2 and R3 (Figure 2—figure supplement 3A-D). In contrast to Kv1.2_s_, neither Shaker and Kv1.2-2.1 structures show this rotation between open and inactivated states (Figure 2—figure supplement 3G-H) but instead the VSDs remain superimposable.

It should be noted that the difference between Kv1.2_s_ and the others in conformational changes might arise from the variety of systems used for structure determination, which include DDM micelles, MSP-bounded nanodiscs, and crystals grown in different laboratories (but under very similar conditions and having the same space group). In the Kv1.2_s_ case for example, the DDM detergent micelles might allow the VSDs to be particularly mobile.

### A dendrotoxin blocks Kv1.2 by inserting a lysine into the pore

Dendrotoxins are peptide neurotoxins from mamba snakes that bind with nanomolar affinities and block potassium channels. Alpha-dendrotoxin (α-DTX) consists of a peptide chain of 59 amino acids stabilized by three disulfide bridges (Fig. 3A) and, like other dendrotoxins, exhibits the same fold as Kunitz protease inhibitors (Skarzynski, 1992). In α-DTX arginine and lysine residues are concentrated near the N-terminus (Arg3, Arg4, Lys5), the C-terminus (Arg54, Arg55) and at the narrow β-turn region (Lys28, Lys29, Lys30). The Lys5 side chain protrudes from the surface of the molecule, and its modification by acetylation markedly reduces binding (Harvey et al., 1997). Replacement of Lys5 by alanine or ornithine decreased the affinity for potassium channels by 1000-fold and 100-fold, respectively, suggesting that the sidechain geometry is critical (Gasparini et al., 1998). The Lys5 sidechain therefore has been the prime candidate for the pore-blocking moiety.

**Figure. 3.**
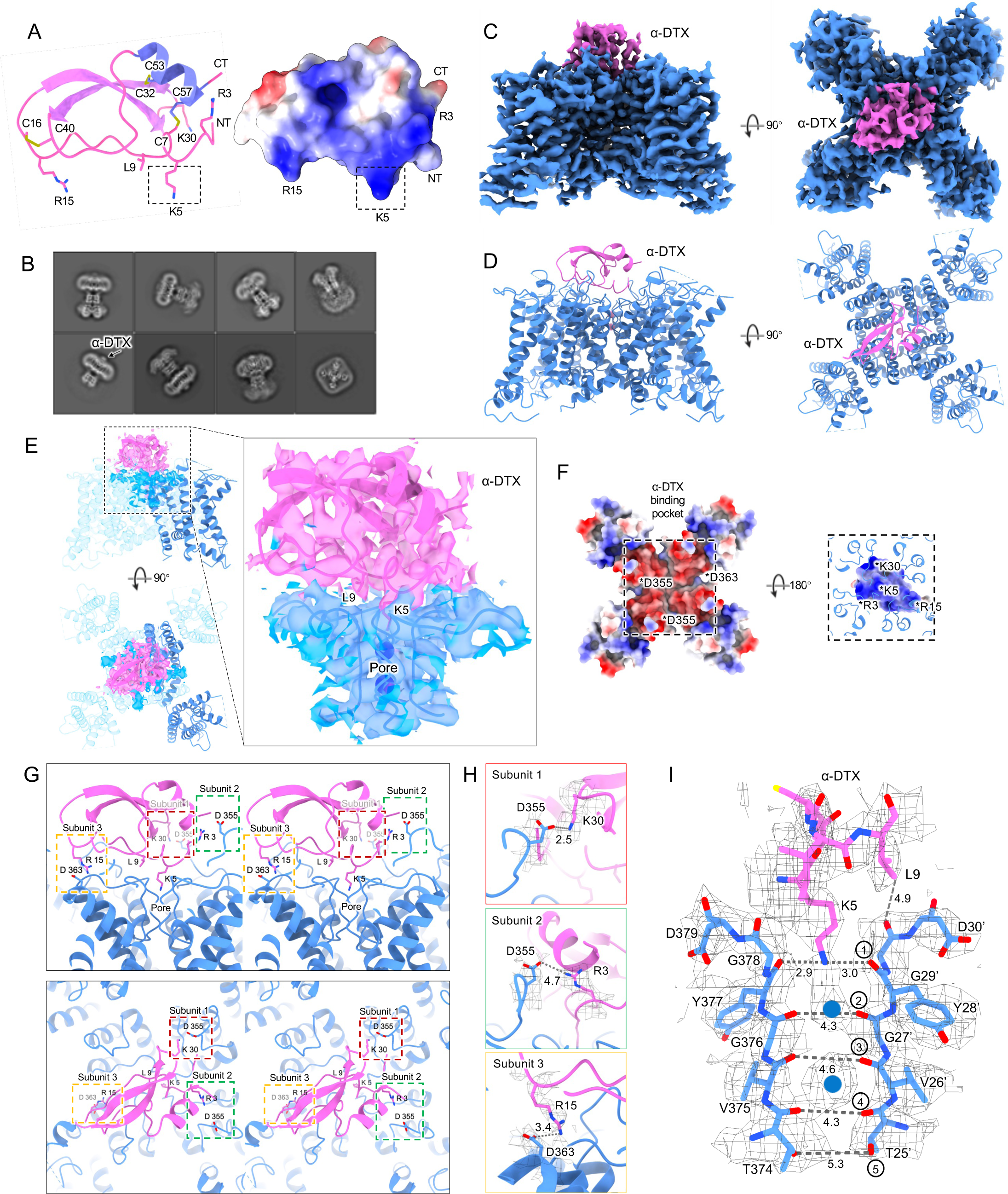
Kv1.2_s_ DTx-bound structure. (A) DTx structure and electrostatic surface view. Basic residue sidechains are illustrated, and disulfide bridges are shown in green. (B) Representative 2D classes from the DTx-bound Kv1.2_s_ . Box size is 273 Å. (C) Side and top view of Kv1.2-DTx cryo-EM density map, obtained with no symmetry imposed. (D) Side and top view of the fitted model. (E) Interaction interface of DTx and Kv1.2_s_, illustrated with map density of the toxin and pore domain, with superimposed models including one complete subunit. (F) Top-down view of the Kv1.2s-DTx structure with DTx removed, showing the channel in electrostatic surface view (left panel); and the corresponding bottom-up view through the DTx-channel interface (right panel) where DTx is shown in electrostatic surface view, and extensions of Kv1.2s helices in blue. The asterisks indicate the charged groups. (G) Three salt-bridge interactions between DTX and Kv1.2_s_ are shown as side (upper panel) and top (lower panel) stereo view. (H) Close-up view of the three salt-bridge interactions from the horizontal plane with cryo-EM density map. (I) Side view of the selectivity filter of Kv1.2_s_-DTx shown with the toxin model for residues K5 to L9. Dashed lines show approximate distances between carbonyl oxygens. Potassium ions are shown as blue balls.

By adding an excess of α-DTx to detergent-solubilized Kv1.2 protein we obtained a dataset of about 300,000 particles where an additional cap of density can be seen outside the pore (Figure 3B). In 3D classification only a small fraction (<1%) of toxin-free particles was detected, and this class was removed before further processing. The processing of particle images is summarized in Figure 3 – figure supplement 1. Refined structures with imposition of four-fold symmetry (C4) or with no symmetry (C1), yielded resolutions 2.8 and 3.2 Å respectively. In the C4 reconstruction the additional cap of density from the toxin was not interpretable, as it arises from the four-fold superposition of the asymmetric toxin bound in alternative poses to the symmetric channel. Zhang et al. (Zhang et al., 2021) have demonstrated however that small asymmetric ligands can be distinguished in large single-particle datasets by focused classification and refinement. Similarly we used symmetry expansion, a reference mask covering only the toxin and upper selectivity filter, and a set of four asymmetric starting references to obtain, through 3D classification without alignment in the Relion software, a C1 reconstruction. The refined C1 density map and model are shown in Figure 3C, D. Due to the C4 symmetry of Kv1.2, there are four equivalent DTx binding sites on the channel. The top and side views display the DTx on the negatively charged outer mouth of the Kv1.2 channel pore, with the density map fitted to the atomic model (Figure 3E).

The X-ray structure of α-DTX (Skarzynski, 1992) could be docked with minor adjustments into the map, with the Lys5 sidechain extending into the channel pore. The positively-charged toxin is tethered to the negatively-charged outer mouth of the pore by three salt bridges: toxin residues K30, R15 and R3 bind to three channel residues D355, D363 and D363 (D6’, D14’ and D14’ in the turret and pore helix). Each of the three channel residues is located on a different subunit (Figure 3G, H).

In the selectivity filter of the toxin-bound channel (Figure 3I) a continuous density is seen to extend downward from the toxin to the boundary of IS1. This density is well modeled by the side chain of Lys5, with the terminal amine coordinated by the carbonyls of Y377 (Y28’). We conclude that block by α-DTx is occlusion of the selectivity filter through binding of the Lys5 terminal amine. In the selectivity filter a large density peak, presumably a K^+^ ion, is seen in a somewhat offset IS1 binding site; a second, larger K^+^ ion peak is seen in IS3.

Like the nearby Lys5 residue, substitution of alanine for Leu9 also results in a 1000-fold reduction in toxin affinity (Gasparini et al., 1998). In the structure, however, the Leu9 sidechain does not show contacts with either the channel or other residues in the toxin itself. It is therefore not obvious why it is important in toxin binding.

We also tried mixing an excess of α-DTX with the Kv1.2s W366F protein sample in an attempt to observe a pore-blocker-bound inactivated structure. However, we found no evidence of bound toxin density on the channel in the cryo-EM 2D classes. This is no surprise in view of the large rearrangement of the extracellular pore entrance in the inactivated state, where the IS1 and IS2 sites are abolished and the corresponding carbonyls point away from the pore axis (Figure 2E), eliminating the site of binding of the lysine side chain. It should be noted however that in the case of a ShK toxin derivative binding to the Kv1.3 channel, Tyagi et al. (2022) observed that binding occurs to inactivated channels. In that case the inactivated state has a less drastically remodeled selectivity filter, and binding of the toxin restores the selectivity filter to its conducting configuration (Tyagi et al., 2022).

### The Kv1.2 selectivity filter in nominally K^+^-free medium

The X-ray crystal structure of the KcsA channel in low K^+^ solution (Zhou et al., 2001) shows a collapsed selectivity filter, in which the residues V26’ and G27’ are rearranged, abolishing the binding sites IS2 and IS3. In low K^+^ other channels including KirBac2.2 and K2p10.1 show large conformational fluctuations of the outer pore in low K+ (Matamoros et al., 2023; Wang et al., 2019), and in the model potassium channel NaK2K the outer half of the selectivity filter unravels at low K^+^ (Sauer et al., 2011). On the other hand, Shaker channels are seen to conduct Na^+^ in the absence of K^+^ (Melishchuk et al., 1998). Thus it is of interest to observe the structure of Kv1.2 under this condition.

We obtained the structure of Kv1.2_s_ in a nominally zero K^+^ solution, with potassium replaced with sodium in all buffers from affinity purification to size-exclusion chromatography (see Methods). We were surprised to find that it is little changed from the K^+^ bound structure, with an essentially identical selectivity filter conformation (Figure 4B and Figure 4—figure supplement 1). Unfortunately, Na^+^ and K^+^ ions cannot be distinguished in the density maps, and we therefore cannot rule out the possibility that peaks in the selectivity filter region might come from contaminating K^+^ ions. Like Kv1.2_s_ in the usual potassium solution, the major density peaks are seen in the IS1 and IS3 ion binding sites, with somewhat lower occupancy of IS2 and IS4 (Figure 4B and E). Increased asymmetry in ion occupancy is expected to accompany a reduction in ion conductance (Morais-Cabral et al. 2001).

**Figure. 4.**
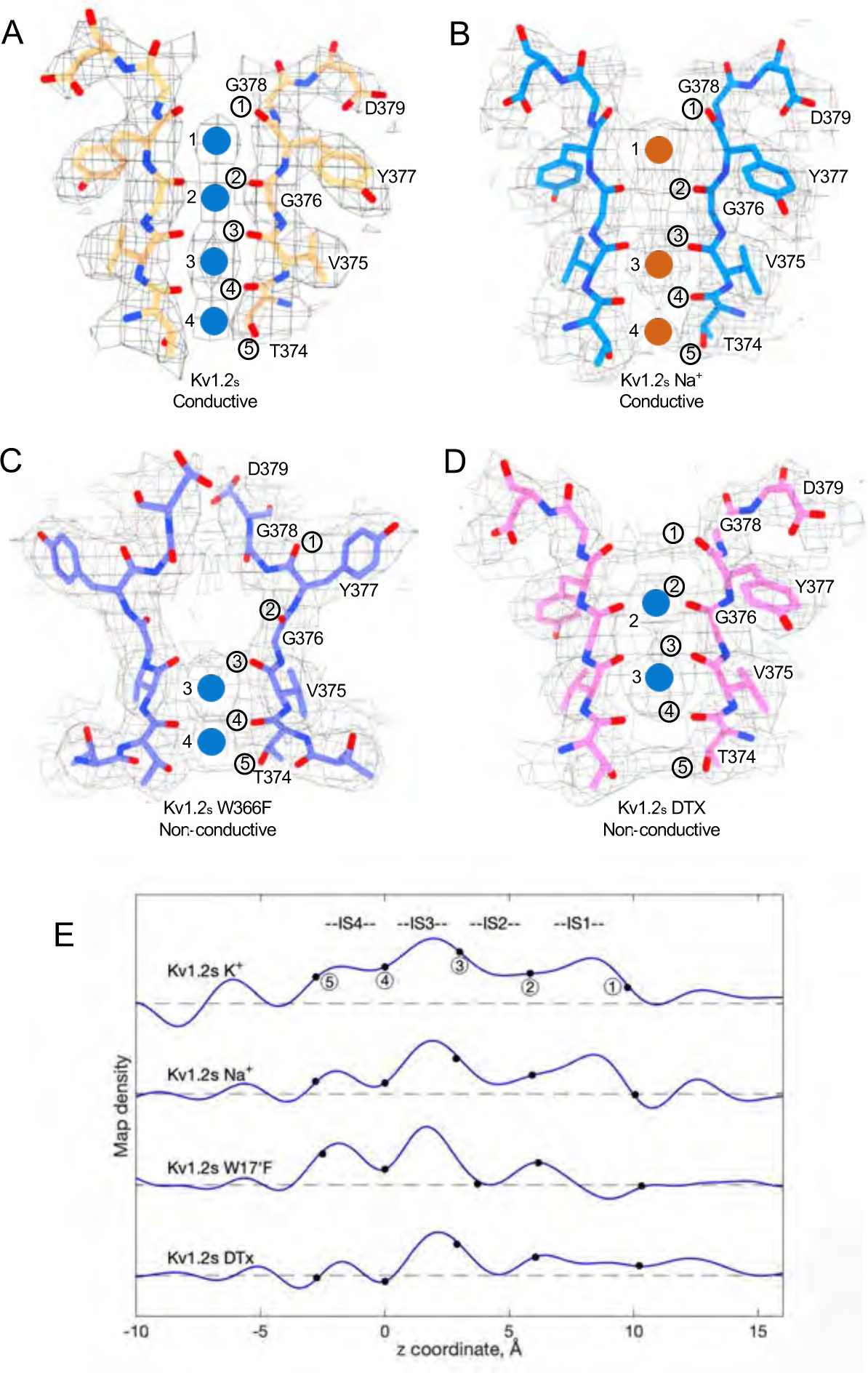
Summary of Kv1.2 conductive and non-conductive pores. Selectivity filter structures of (A) Kv1.2_s_, (B) Kv1.2_s_ Na^+^-bound, (C) Kv1.2_s_ W366F, and (D) Kv1.2_s_-DTX. Opposing pairs of subunits are shown, and potassium ions and sodium ions are shown as blue and orange balls, respectively. Circled numbers label the P-loop ion-coordinating oxygens. (E) Plots of density along the symmetry axis of the selectivity filter region. Values of the z-coordinate are relative to the position of the T25’ (T374) carbonyl oxygen. The nominal positions of the coordinating oxygens are marked and numbered as in parts A-D. Dashed baselines indicate the external solvent density. The scaling of the density traces is arbitrary. The DTx map was computed with no symmetry imposed; for the others, C4 symmetry was imposed in reconstruction.

We conclude that the Kv1.2s selectivity filter is stable in low K^+^ solutions, and that the channel most likely conducts Na^+^ in the absence of potassium; both of these features have been observed in the NaK2K channel (Sauer et al., 2013), where the electron density profile in the selectivity filter in Na^+^ is little changed when K^+^ is added in a small amount—1 mM—that produces partial K^+^ occupancy. Na^+^ conduction is also observed in the non-inactivating KcsA E71A channel (Mita et al., 2021).

### Large conformational fluctuations of the W366F mutant in K^+^-free medium

We also collected cryo-EM data from the Kv1.2_s_W17’F mutant channels, using the same procedures to replace K^+^ with Na^+^ during purification. The corresponding Shaker-W17’F channel stably carries large Na^+^ and Li^+^ currents in the absence of K^+^ (Starkus et al., 1998). In 2D classification of our single-particle images we saw large variability in the structure, as if the connection between the transmembrane domain and the intracellular domains (the T1 domain and the Kvβ2 subunits) was highly flexible (Figure 5A-B). The intracellular domains were resolved to 3.0 Å resolution in a focused reconstruction (Figure 5C) but the complementary focused reconstruction of the transmembrane domain yielded only about 7 Å resolution. Nevertheless this low-resolution map matches the secondary structure of the Kv1.2_s_ W366F mutant in K^+^ solution (Figure 5D), with the possible exception of low density in the selectivity filter region (Figure 5E-F). We conclude that the structure of this mutant channel becomes unstable in the absence of K^+^, as if the tight binding of K^+^ ions is required to stabilize the altered selectivity filter.

**Figure 5.**
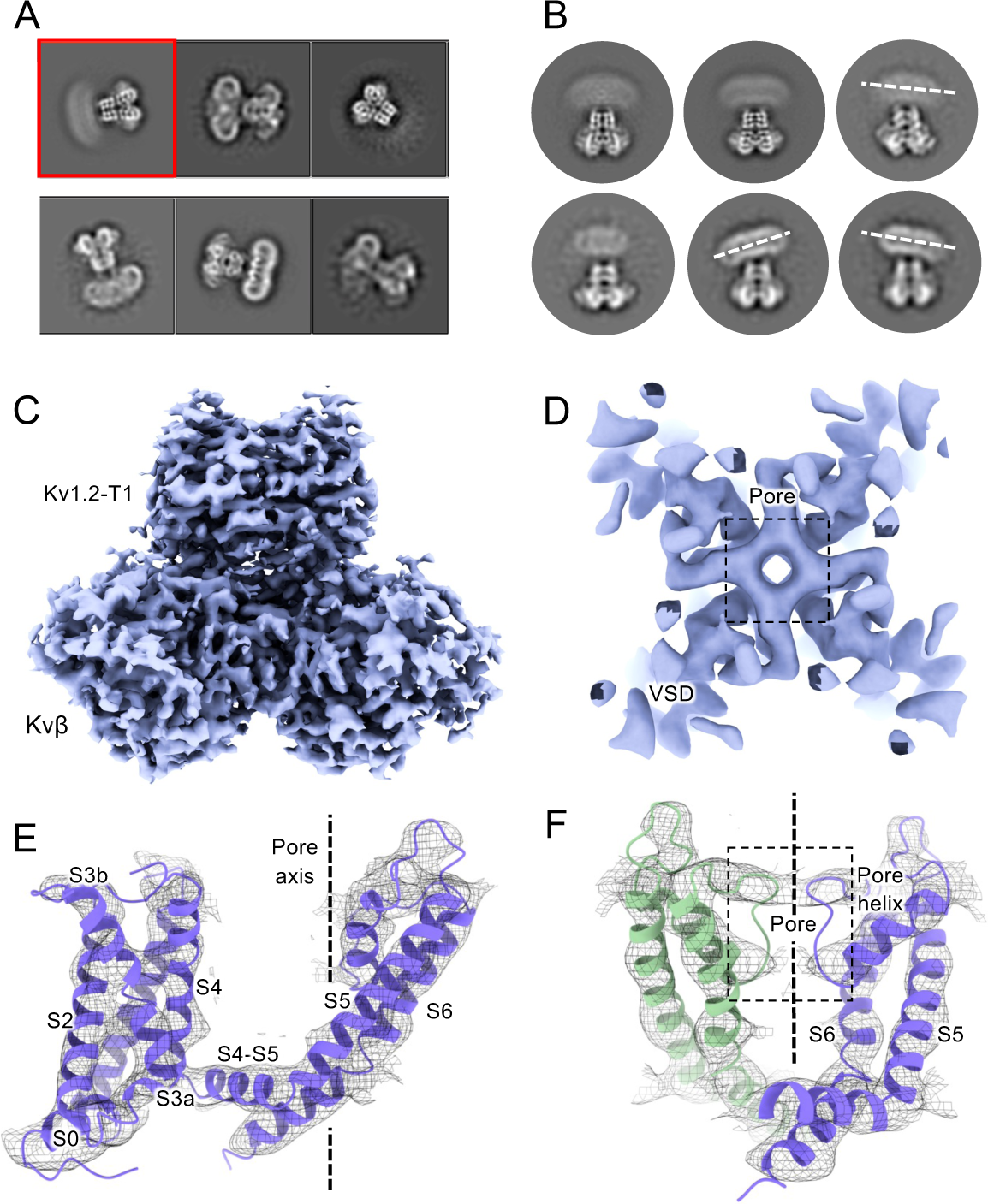
Cryo-EM analysis of Kv1.2 W366F in Na^+^. (A) Representative 2D classes. (B) Second round of classification of first class in (A), outlined in red. Wobbling of the transmembrane domain is illustrated by the white dashed lines. (C) Density of intracellular domain, obtained by focused refinement. (D) Top view of TMD reconstruction, with resolution 7.0Å. (E) Overall TMD map density fits with the Kv1.2 W366F model, shown as ribbons. (F) There is low density in the selectivity filter (dashed rectangle).

## Discussion

The potassium channel signature sequence TTVGYGD was first identified by Heginbotham et al. (Heginbotham et al., 1994) as critical for potassium selectivity. The residues fold as an extended loop (the P-loop) and is part of the P-region, about 20 residues of conserved sequence located between the S5 and S6 helices. We use here a system of numbering the entire linking region between S5 and S6, as identified in Shaker by Miller (Miller, 1990). The P-loop (residues 24’-30’ in this numbering, TTVGYGD) forms four ion binding sites (Doyle et al., 1998) through which K^+^ ions and water molecules can pass sequentially in a knock-on fashion. The geometry and flexibility is finely tuned to allow high K^+^ selectivity with a high transport rate (Noskov & Roux, 2006; Roux, 2005).

In this paper we have considered the ion occupancy and conformation of the P-region under four conditions: the conducting state, a C-type inactivated state, the channel blocked by dendrotoxin, and in the absence of permeant ions. In the conducting state, the cryo-EM structure of our rat Kv1.2 construct (Kv1.2_s_) in DDM detergent micelles agrees with the crystal structure of Long et al. (2005) and is very similar to the structure of *Drosophila* Shaker in lipid nanodiscs (Tan et al., 2022). Also, apart from local differences in the chimeric region of the voltage sensor domains (VSDs), we confirm the observation (Long et al., 2007) that the rat Kv1.2 structure is essentially identical to the structure of the rat Kv1.2-2.1 paddle chimera as obtained from X-ray crystallography. In turn, the structure from cryo-EM imaging of Kv1.2-2.1 channels in lipid nanodiscs (Matthies et al., 2018) is also nearly identical to the X-ray structure. Thus the conformation of the entire transmembrane region of the Kv1.2 channel and its variants appears to be remarkably insensitive to its environment, whether lipid or detergent.

### Inactivation

After the opening of the intracellular activation gate formed by the S6 helices, C-type inactivation (Hoshi et al., 1991) causes channel currents to switch off spontaneously. The inactivation process is very sensitive to mutations in the selectivity filter and nearby residues of the P-region and is influenced by the state of the S6 gate (Cuello et al., 2017; Li, Shen, Rohaim, et al., 2021) and the VSDs (Bassetto et al., 2021; Conti et al., 2016). Low K^+^ concentrations, especially in the extracellular solution, accelerate inactivation (Levy & Deutsch, 1996).

Is C-type inactivation a reflection of selectivity filter instability? Sauer et al. (2011) investigated the structural stability of a channel selectivity filter, using a model system based on the NaK prokaryotic nonselective channel (Shi et al., 2006). NaK’s selectivity filter (P-loop sequence TVGDGN) has only two ion binding sites, and opens into a wide extracellular vestibule. A similarly wide vestibule is seen in the nonselective HCN1 channel (Lee & MacKinnon, 2017; Shi et al., 2006) (Figure 6). NaK can however be converted to a highly K^+^ selective channel, termed NaK2K, through just the two mutations D28’Y and N30’D, yielding the P-loop sequence TVGYGD. NaK2K has the four-site selectivity filter seen in all K^+^-selective channels, and the residues Y28’ and D30’ participate in a hydrogen-bonded network to stabilize the selectivity filter just as in KcsA channels (Doyle et al., 1998). Y28’ also participates in hydrophobic packing interaction. The removal of any of these interactions results in a loss of selectivity, and unraveling of the selectivity filter, returning to the large vestibule seen in NaK channels. Sauer et al. (2011) conclude that the standard four-ion-site, K^+^ selective configuration of potassium channel selectivity filters is an energetically unfavorable, strained backbone conformation, and the weakening of the specific interactions that hold it in place lead to its unraveling, producing a lower-energy configuration that yields a nonselective conduction path. Similarly, the removal of K^+^ ions from NaK2K destabilizes the selectivity filter, resulting in large fluctuations of the outer pore (Matamoros et al., 2023).

**Figure 6.**
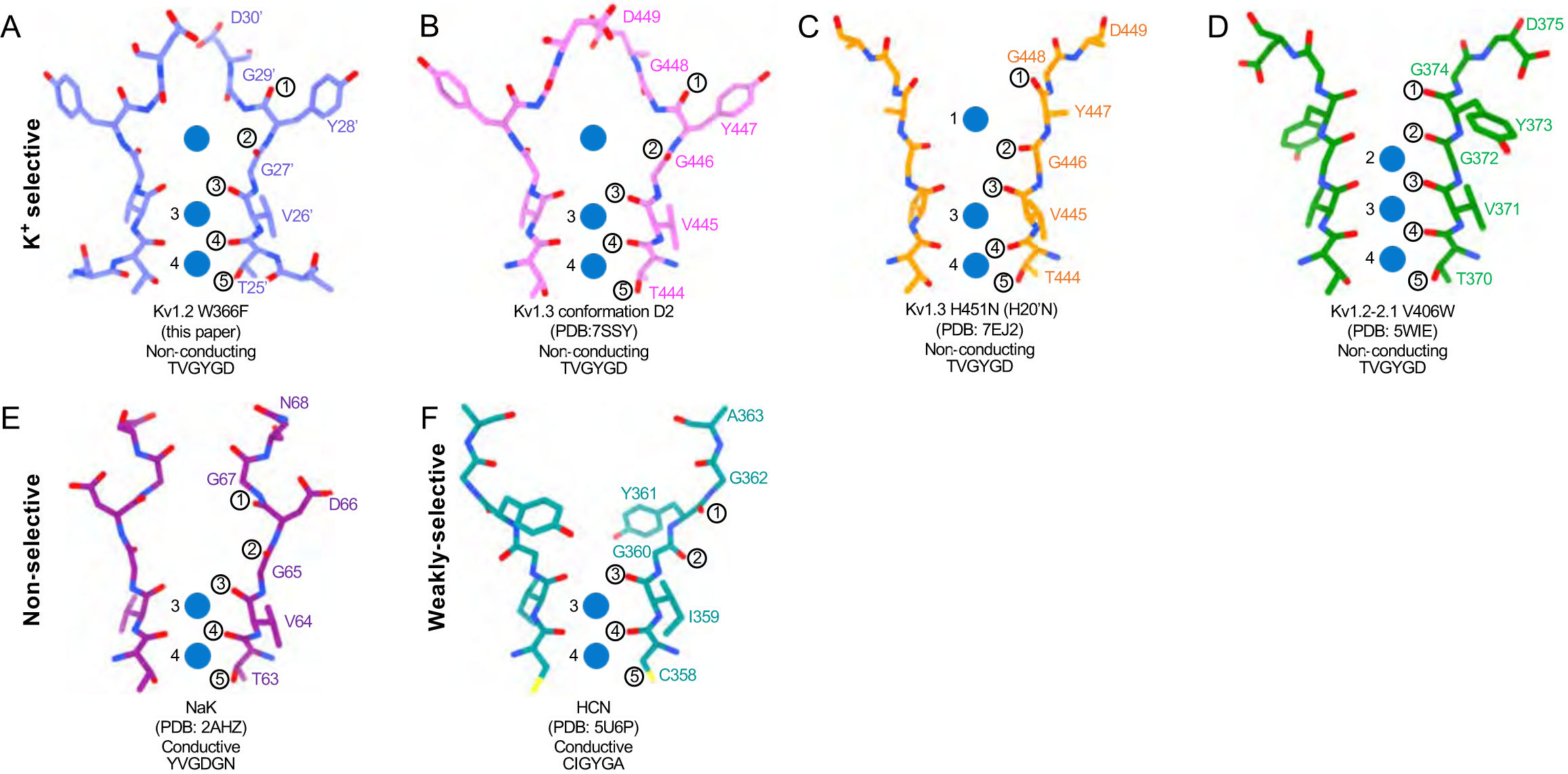
Structural comparison of Kv1.2 W366F with various channel pores. A-D, Selectivity filter structures and P-loop sequences of potassium selective non-conducting channels: (A) Kv1.2_s_ W366F, (B) Kv1.3 alternate conformation, (C) Kv1.3 H451N, (D) Kv1.2-2.1 V406W. E-F, Pore regions of less-selective channels: the non-selective, conducting NaK (E) and the weakly-selective, conducting HCN (F). Potassium ions are shown as blue balls. Circled numbers enumerate the pore-forming carbonyl oxygens; carbonyl 2 faces the adjacent subunit in the clockwise direction in all but the HCN channel in panel F.

The H-bond interactions involving Y28’ and D30’ are also key to C-type inactivation in Shaker channels. Weakening those interactions produce channels that inactivate quickly or are found permanently inactivated (Pless et al., 2013). It is therefore no surprise that the loosening of H-bond restraints, resulting in the unraveling of the selectivity filter, yields a preferred, low-energy conformation with a large extracellular vestibule for the inactivated state of Kv channels.

In the case of Kv1.2, structural and molecular-dynamics studies have focused on the mutation W17’F (W366F in Kv1.2) which induces an inactivation-like process that is slowed in the presence of extracellular K^+^ ions or pore blockers. In Shaker this mutation (W434F) causes the near total loss of open-channel current (Li et al., 2018; Perozo et al., 1993; Yang et al., 1997); in Kv1.2 it greatly accelerates inactivation (Suarez-Delgado et al., 2020). A large expansion of the outer selectivity filter in W17’F mutants is seen in structures of Shaker (Tan et al., 2022), Kv1.2-2.1 channels (Reddi et al., 2022) and also in Kv1.2 as reported here. Quantified as the distance between the diagonally-positioned Y28’ alpha-carbons, the expansion in Shaker is from 8.6 to 12.8 Å. In both the Kv1.2-2.1 paddle chimera (Reddi et al., 2022) and in Kv1.2_s_ the expansion is approximately from 7.2 to 10.7 Å. The expanded outer pore is very similar to the large extracellular vestibules observed in NaK and HCN1 channels (Fig. 7).

A striking feature of the inactivated state structure reported by Tan et al. (Reddi et al., 2022; Tan et al., 2022) is the high occupancy by potassium ions of the two remaining binding sites IS3 and IS4. During normal conduction, ion binding sites in the selectivity filter are usually occupied by K^+^ and water molecules in alternation (Morais-Cabral et al., 2001). However, in molecular-dynamics simulations based on their Shaker W434F structure, Tan et al. (Tan et al., 2022) observed with high probability the simultaneous occupancy of IS3 and IS4 by two K^+^ ions. The ions did not leave the sites during a (relatively long) 2 µs simulation with 300mV membrane potential applied, but knock-on permeation events were observed at a low rate, two events during 1µs, at the very high potential of 450 mV. A similar result was observed in earlier MD simulations of the Kv1.2-2.1 chimera where the W17’F mutation was artificially introduced (Conti et al., 2016; Kondo et al., 2018). Those simulations did not show the large rearrangement of the outer pore, but they did demonstrate small shifts in the geometry of IS3 and IS4 that, like those seen by Tan et al., produced simultaneous occupancy by K^+^ ions and blockage of ion permeation. We also see strong ion densities in the two remaining ion binding sites in the Kv1.2_s_-W17’F channel (Fig. 2E, Fig. 4E), and a similar tight binding of adjacent K^+^ ions seems also to block K^+^ permeation in the structure of an inactivated Kv1.3 channel (Liu et al., 2021).

In a study of the Shaker mutant W17’F, Yang et al. (1997) observed, with symmetrical high K+ concentrations to maximize channel activity, brief (1 ms) openings of channels that occurred with a peak open probability of 10^-5^ immediately following a large depolarization to +80 mV (Yang et al., 1997). When open, the single-channel currents were the same magnitude as the currents in wildtype channels. It is unlikely that these currents arose from rare ion-conduction events through destabilization of the tightly-bound K^+^ ions in the inactivated configuration, as such currents would be too small to be observed. Instead, the simplest explanation is that these currents arise when structural fluctuations transiently reconstitute the normal four-site selectivity filter, and the channel conducts normally for brief intervals. The normal selectivity filter is expected to be stabilized by bound K^+^ ions. In the absence of external potassium ions the peak open probability drops to about 10^-7^ (Yang et al., 1997).

Inactivation is voltage-dependent: it occurs less in channels held at a negative membrane potential, but is enhanced in a time-dependent manner with depolarization. This was observed for example in the experiments of Yang et al. (1997) in our recordings from Kv1.2_s_ W17’F channels (Fig. 1, figure supplement 1). This voltage dependence could arise from a coupling between voltage sensors and the pore domain, which would be reflected in a difference in gating-charge movement (reporting VSD conformational changes) between wildtype and W17’F channels. There is however little difference in gating-charge movement between Shaker and Shaker W17’F channels, although Shaker with certain mutations in S4 and S6 do show differences (Bassetto et al., 2021; Conti et al., 2016). A more important mechanism seems to be coupling between the S6 activation gate of the channel and the inactivation mechanism, such that steps in the physical opening of the channel would in increase the inactivation rate. Such a coupling is clearly seen in a series of KcsA X-ray structures by Cuello et al., (2010): with increased opening of inner bundle gate there is an increased steric clash between residues of the M2 helix and residues on the pore helix that could lead to inactivation. They identified a similar interface in the Kv1.2-2.1 structure, which provides a simple explanation for coupling between channel opening and inactivation processes.

Are the structures of the various W17’F channels valid models for C-type inactivation? In recent cryo-EM studies (Selvakumar et al., 2022) the human Kv1.3 channel was found to spontaneously exist in one or two alternative conformations that appear to be inactivated states. The D1 conformation shows a large displacement of D30’ and is reminiscent of the cryo-EM structure of the Kv1.3 rapidly-inactivating mutant H20’N (Liu et al., 2021). It exhibits a partially twisted P-loop (Figure 6 and Figure 6—figure supplement 2B) as if it were an intermediate state between open and the W17’F structures. Meanwhile the D2 conformation is very similar to that of the W17’F variants considered here. This strongly supports the widely-held view that these mutant channels are in a conformation close to, if not identical to, a true C-type inactivated state.

Why is inactivation much more complete in the Shaker mutant W17’F than in Kv1.2 or Kv1.2-2.1? Clearly one can compare the stabilities of the conducting and the inactivated conformations. Wu et al. (2022) mutated several of the residues that form important interactions in these channels, and found that the inactivation behaviors of Kv1.2 and Kv1.2-2.1 are different from that of Shaker. In the conducting state, the H-bond networks stabilizing the key residues Y28’ and D30’ in Shaker and Kv1.2 very similar, with the only obvious difference being at position 22’ where Shaker has Thr but Kv1.2 has Ser. It turns out that the mutation S22’T in the Kv1.2 background has little effect on inactivation; thus to a first approximation the conducting state is equivalently stabilized.

In the inactivated state structures, all obtained with the W17’F mutation, a variety of H-bond interactions arise in the linker sequence between the P-loop and the S6 helix, with the variety arising from different conformations of Loop 1 and Loop 2, two regions of the linker (Figure 6—figure supplement 1). Details of the interactions are discussed in the Figure legend. We conclude that a quantitative evaluation of the stability of the inactivated state will require more than simply an enumeration of H-bonds in this region of the protein sequence.

### Block of the channel by dendrotoxin

Potassium currents can be inhibited by five distinct families of toxins. The spider toxins Hanatoxin, SGTx1 and VSTx1 interact with the voltage sensors of Kv channels to inhibit voltage-dependent channel activation (Lee & MacKinnon, 2004; Wang et al., 2004). Other toxins bind directly to the extracellular mouth of the channel to block current. The cone-snail toxin Cs1 binds to the extracellular surface of Kv channels and is thought to block currents by inducing a conformational change in the protein, collapsing the selectivity filter (Karbat et al., 2019). In X-ray crystal structures the scorpion toxin Charybdotoxin is seen to block Kv1.2-paddle chimera channels by inserting a lysine residue directly into the selectivity filter, disrupting the ion-binding site IS1 to inhibit ion flow (Banerjee et al., 2013). The same mechanism holds for block of Kv1.3 by the sea anemone toxin ShK as studied by cryo-EM (Selvakumar et al., 2022); in the latter two cases the lysine terminal amine interacts with the Y17’ carbonyls which form the top of binding site IS1. We now report that Kv1.2 block by the snake toxin α-Dendrotoxin (α-DTx) is by a similar mechanism.

Obtaining the structure of an asymmetric ligand bound to a four-fold symmetric channel, is challenging, and the four structural studies that have been done have used different strategies. In their X-ray crystallographic study, Banerjee et al. (2013) were able to interpret and model at low resolution the four-fold superimposed toxin density, using heavy-atom derivatives in a series of crystal structures to establish the pose of the toxin molecule. The difficulty of modeling the CTx molecule precluded assigning a precise location to the Lys22 channel-blocking sidechain; nevertheless, in the 2.5 Å map of the selectivity filter are seen three well-resolved, discrete ion densities in the sites IS2, IS3 and IS4, and the authors conclude that the toxin interaction prevents ion occupancy of the IS1 site. A cryo-EM study that yielded a similar C4 symmetric reconstruction of Kv1.3 with bound dalazatide, a derivative of the ShK toxin, was obtained by Tyagi et al. (2022). The C4-symmetric reconstruction did not yield an interpretable map of the toxin, but clear ion densities in IS2, IS3 and IS4 were visible.

Selvakumar et al. (2022) also obtained a structure of ShK bound to Kv1.3, but in this case the ShK toxin was fused to a Fab fragment; this yielded an asymmetric ligand large enough to allow C1 reconstruction from the cryo-EM images. The resulting 3.4Å map allowed the toxin K22 sidechain to be modeled entering the pore and extending to near the Y17’ carbonyl oxygens.

From our cryo-EM data we obtained a C1 map of α-DTx bound to Kv1.2_s_ at 3.2 Å; this was done by extensive image processing and was helped by the larger size of α-DTx (59 residues as compared to 35 for ShK). With this map the side chain of the toxin residue K5 could be traced, apparently with the terminal amine directly coordinated by the Y17’ carbonyls. A surprise in this map are the positions of the K^+^ ion peaks in the selectivity filter (Figure 4E). Ion density is visible in IS3, and somewhat lower density is seen in IS1; there is no density peak in IS2. The different pattern of ion densities with DTx compared to CTx, having two rather than three peaks in the selectivity filter, implies that the interaction between the lysine side chain and the selectivity filter is in some way different in DTx compared to the other toxins.

### The selectivity filter in the absence of K^+^

In the absence of K^+^, large voltage-dependent Na^+^ currents are observed in wildtype Kv2.1 channels (Korn & Ikeda, 1995) and in Shaker channels (Melishchuk et al., 1998). The Kv1.2 structure reported here with Na^+^ replacing essentially all K^+^ in the solution shows a selectivity filter conformation that is little changed from the K^+^ structure (Figure 5B). Ion densities are seen in the ion-binding sites, and the occupancies of the sites are altered, but the limited resolution precludes an analysis of the coordination of ions.

The Shaker-W17’F channel in the absence of K^+^ stably carries large Na^+^ and Li^+^ currents (Starkus et al., 1997). These currents are blocked by low concentrations of K^+^, and this would be expected as potassium binding to the sites IS3 and IS4 in this mutant is very tight (Tan et al., 2022). We therefore sought to obtain the structure of the corresponding Kv1.2-W17’F channel in Na^+^ solution (Figure 5). In 2D classes the complex appeared to be highly unstable, with large variations in the “wobble” angle between the transmembrane and cytoplasmic domains. Attempts to recover the structure of the transmembrane region by focused refinement yielded only low-resolution structural information. We conclude that this channel, at least as solubilized in detergent, has a highly unstable conformation when K^+^ ions are absent.

## Methods

### Kv1.2-Kvβ expression and purification

The rat Kv1.2 alpha-subunit constructs were derived from that of Long et al. (2005), having the full-length (GenBank: X16003) sequence containing the mutation N207Q to eliminate a glycosylation site. Our Kv1.2_s_ construct contained the additional mutations L15H and G198S in unstructured regions; the inactivating construct contained the further mutation W366F. To each of these was added at the N-terminus two tandem Strep tags (sequence WSHPQFEK) separated by a Gly-Ser linker. The beta-subunit construct was derived from the rat beta2 core, residues 36-357 (Gulbis et al., 1999). To eliminate very strong electrostatic interactions between the many Lys residues on the “bottom” of the beta-subunits and the negatively-charged carbon film or graphene substrates, we mutated to glutamine the five lysine residues at positions 94, 104-106 and 258 of the beta2 subunit. The fold of the beta subunits containing these mutations is indistinguishable from the wildtype structure of Long et al. (2007).

To express both subunits in a single construct, Kv1.2 and Kvβ genes were separately inserted into pPicZ-B (Thermo-Fisher) vectors between the XhoI and AgeI sites in the multi-restriction-sites region. Subsequently the pPicZ-B vector carrying Kvβ was opened at BglII and BamHI restriction sites, and the insert including the AOX1 promoter was inserted into the other pPicZ-B vector, upstream of the AOX1-driven Kv1.2 cDNA, at the BglII site.

The *P. pastoris* strain SMD1168 (Invitrogen) was electroporated with the pPicZ-B plasmids. YPDS (yeast extract, peptone, dextrose, and sorbitol) plates containing Zeocin (800 ug/ml) were used to select transformants. Stocks were stored in 15% glycerol at -80°C.

The expression and purification procedures were based on those of Long et al. (Long et al., 2005). Briefly, we inoculated 1 liter of BMGY medium (this and the other yeast media were obtained from Invitrogen) with 10 ml of an overnight culture grown in YPD medium with Zeocin (100 ug/ml). After centrifugation (5500× %, 10 minutes), the cells were transferred to BMMY medium including 0.5% (v/v) methanol and grown for 24 h. After adding an additional 0.5% methanol to increase protein expression, cells were grown for 2-3 days. Cells were centrifuged (5500× %, 20 minutes), the pellet was frozen and stored at -80°C.

Thawed cells were lysed at 4° in a French Press. One gram of cell lysate was resuspended in 5mL of Lysis buffer consists of 100 mM Tris-HCl (pH 8.0), 150 mM KCl, 300 mM sucrose, 5 mM EDTA, 0.05 mg deoxyribonuclease I, 1 mM MgCl_2_, 1 mM phenylmethylsulfonyl fluoride, 1 mg/ml each of leupeptin, pepstatin and aprotin, and 0.1 mg/mL of soy trypsin inhibitor. Lysis buffer plus 30 mM dodecylmaltoside (DDM) was used to solubilize the membranes for 3 hours at room temperature. Centrifugation (13000×*g*, 20 minutes) separated the unsolubilized material. The supernatant was added to Strep-Tactin Sepharose beads (IBA Lifesciences) preequilibrated with Membrane Resuspension Buffer (100 mM Tris-HCl pH 8.0, 150 mM KCl, 3mM TCEP, 5 mM EDTA, 10 mM beta-mercaptoethanol, and 5 mM DDM. The bead slurry (approximately 1 ml) was incubated 1h at 4°C with gentle rotation. A column was used to collect the beads after incubation, followed by washing with 14 volumes of Wash Buffer, which was Membrane Resuspension Buffer with added lipids (0.1 mg/ml of the mixture 3:1:1 of POPC:POPE:POPG). The addition of 10 mM desthiobiotin was used to elute the bound protein. The eluted protein was concentrated with a Millipore Amicon Ultra 100 K filter and further purified by size-exclusion chromatography (SEC) on a Superose S6 column pre-equilibrated with Running Buffer: 20 mM tris-HCl (pH 7.5), 150 mM KCl, 2 mM TCEP, 10 mM DTT, 1 mM EDTA, 1 mM PMSF, 1 mM dodecylmaltoside (Anatrace, anagrade) and 0.1 mg/ml lipid mixture (3:1:1 of POPC:POPE:POPG). We pooled fractions containing both alpha and beta subunits, and when necessary concentrated the protein to 1∼2 mg/ml (Amicon Ultra 100 KDa, Millipore). For experiments under K^+^-free conditions, the protein bound to Strep-Tactin was washed with 5 column volumes of K^+^-free Wash Buffer (150 mM NaCl replacing KCl) and eluted in 3 mL of K^+^-free elution buffer. As for SEC, the column was first washed with three volumes of K^+^-free Running Buffer before running the protein sample.

### Oocyte expression and voltage-clamp recordings

The Kv1.2_s_ alpha subunit or its W366F mutant construct was cloned into a pcDNA3.1 vector. RNA was prepared from the XbaI-linearized plasmid using T7 RNA polymerase. Xenopus oocytes were defolliculated by collagenase treatment, injected with cRNA and stored in ND96 solution (96 mM NaCl, 2 mM KCl, 1.8 mM CaCl2, 1 mM MgCl_2_, 5 mM HEPES, pH 7.4 with NaOH) at 18°C. Recordings were done at room temperature, 5-6 days post-injection in ND96 solution or KD96, the same solution but made with 96 mM KCl, 2 mM NaCl and KOH. Two-electrode voltage clamp recordings employed an OC-725C amplifier (Warner Instruments).

### Cryo-EM specimen preparation, data acquisition and processing

Quantifoil holey carbon grids (R1.2/1.3, 300 mesh, Au) were glow-discharged at 15 mA for 1 min with carbon side facing upwards in the chamber. The chamber of the freezing apparatus (Vitrobot Mark IV, Thermo Fisher Scientific) was preequilibrated at 16 °C and 100% humidity. A 3 μL droplet of protein sample was applied to the carbon side of each grid and blotted for 3 or 5 s with zero blotting force after 15 s wait time before plunge-freezing in liquid ethane. For the DTX bound sample, about 400nM α-DTX (Alomone Labs) was mixed with Kv1.2_s_ protein 30 min prior to cryo-EM grid preparation.

For the Kv1.2_s_-Na^+^, W366F-Na^+^, and DTX-bound samples, graphene-covered grids were used to allow lower protein concentrations (0.05 to 0.1 mg/ml) to be used. These were prepared from commercial graphene (Trivial Transfer Graphene, single layer; ACS Material LLC) using the recommended transfer protocol and floated onto quantifoil grids. To make the graphene layer hydrophilic, grids were then plasma-cleaned (Gatan 950 Advanced Plasma System) under Ar/O_2_ at 10 W for 10 s with the graphene coated surface facing upwards in the chamber (Fan et al., 2019). Vitrification was conducted as described above but with 30 s wait time.

Cryo-EM samples were imaged on a Titan Krios cryo-EM (Thermo Fisher Scientific) operated at 300 keV, equipped with a Bio Quantum energy filter (20 eV slit width) and K3 camera (Gatan). Micrographs were collected using super-resolution mode, physical pixel size 1.068 Å, with SerialEM software (Schorb et al., 2019) using image shift patterns and one shot per hole. A total of 6335, 8310, 4507, 2572, 4680 micrographs were collected for WT, W366F, DTX-bound, Na^+^ bound WT, and Na^+^ bound W366F, datasets respectively.

Processing for each dataset is summarized in the Figure Supplements to Figs. 1-4. Beam-induced motion was corrected by MotionCor2 (Zheng et al., 2017) and CTF values were estimated by Gctf (Zhang, 2016). Particles were picked either by reference-free Laplacian-of-Gaussian autopicking; an adversarial-template-based program Demo Picker written by F.J.S.; or RELION reference-based auto-picking. All further processing used RELION 3.1. Picked particles were extracted and subjected to rounds of 2D and 3D classification before refinement. Transmembrane-domain masks were used for focused refinement to optimize the resolution of the transmembrane region. Beam-tilt and per-particle CTF refinement followed by per-particle motion correction were applied to further improve the resolution.

Analysis of the DTx-Kv1.2_s_ complex started with a C4 reconstruction of the complex from 323k particles. We then performed a rough docking into the Kv1.2_s_ apo density of a toxin map created from the 1DTX model. The toxin pose was chosen not to be inconsistent with the symmetrized toxin density visible in the DTx-Kv1.2_s_ C4 map. Four of these composite maps were created, with the toxin pose rotated 90° each time about the z-axis. These were filtered to 20Å resolution for use as 3D references. Meanwhile a C4 symmetry expansion of the 323k particles was performed, resulting in a 1.294M particle stack. A soft cylindrical mask of radius 24 Å and height 48 Å was created, centered on the z-axis at the interface between the toxin and the pore region. Using this mask and the four 3D references, we performed 3D classification without alignment in RELION, and chose one of the four resulting classes as a C1 reconstruction of the masked region. Using the corresponding set of 312k particles we performed C1 homogeneous reconstruction in CryoSPARC to yield the refined structure at 3.2 Å resolution.

### Protein model building, refinement and structural analysis

An Alpha-Fold predicted rat Kv1.2 model (alphafold.ebi.ac.uk/entry/Q09081) was used to build the Kv1.2 native atomic model. The W366F, DTX-bound, and Na^+^-bound models were subsequently built from this Kv1.2 model. Model fitting was performed with the CCPEM program suite. Initial docking was performed with Molrep; obvious outliers of were manually fixed in Coot 0.9 (Emsley et al., 2010) and real space refinement used Refmac 5 (Murshudov et al., 2011) and Phenix (Liebschner et al., 2019). Density map rendering, analysis and figure preparation were done with UCSF Chimera and ChimeraX (Goddard et al., 2018; Pettersen et al., 2004).

## Acknowledgements

We thank Marc Llaguno and Jianfeng Lin for aid with microscopy for screening on the Glacios microscopes, and Shenping Wu for aid with the Titan Krios data collection. We are grateful to Yufeng Zhou (University of Pennsylvania) and David Fedida (University of British Columbia) for providing the original constructs and consultation on expression constructs. We are grateful to the Yale Center for Research Computing and to the SBGrid Consortium for software support.

## Author contributions

Yangyu Wu, Data curation, Formal analysis, Investigation, Methodology, Writing - original draft; Yangyang Yan and Youshan Yang, Data curation, Formal analysis, Investigation, Methodology; Shumin Bian, Alberto Rivetta and Ken Allen, Methodology; Fred Sigworth, Conceptualization, Formal analysis, Supervision, Validation, Funding acquisition, Project administration, Writing – review and editing.

## Data availability

The cryo-EM maps and atomic models for this work have been deposited in the Electron Microscopy Data Bank (https://www.ebi.ac.uk/pdbe/emdb/) and in the Protein Data Bank (https://www.rcsb.org). The data are available under the following accession codes: Kv1.2 open (EMD-43134, PDB 8VC6), Kv1.2-W366F inactivated (EMD-43136, PDB 8VCH), Kv1.2-DTx toxin-blocked (EMD-43131, PDB 8VC3) and Kv1.2 in Na^+^ (EMD-43133, PDB 8VC4). The particularly challenging cryo-EM dataset of Kv1.2-W366F in Na^+^ will be deposited in the Electron Microscopy Public Image Archive.

---

**Figure 1 - figure supplement 1.**
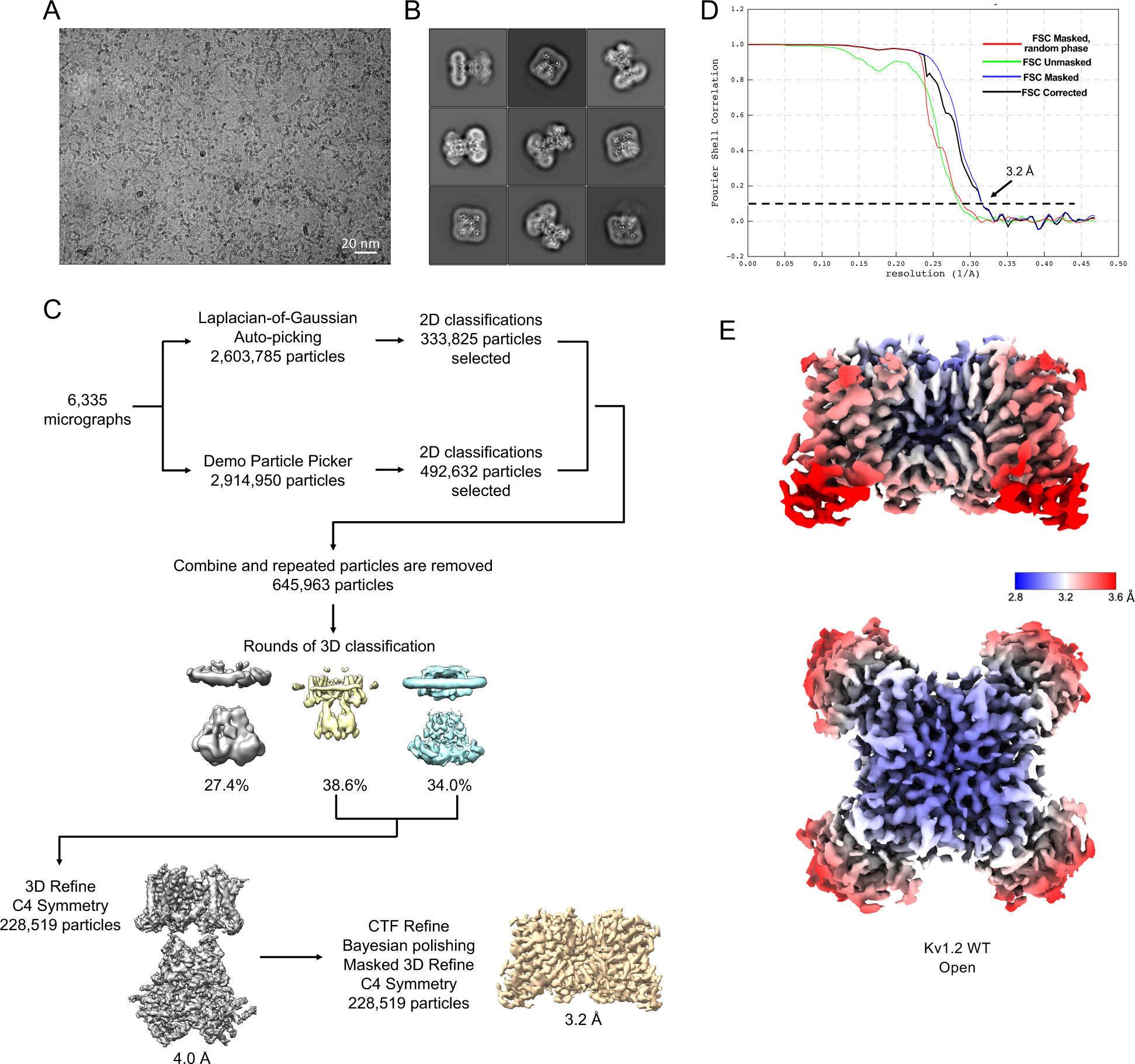
Image processing and reconstruction of Kv1.2_s_. (A) Representative micrograph. (B) Representative 2D classes. (C) Cryo-EM data processing workflow. (D) Gold standard FSC resolution estimation. (F) Local resolution estimation.

**Figure 2 - figure supplement 1.**
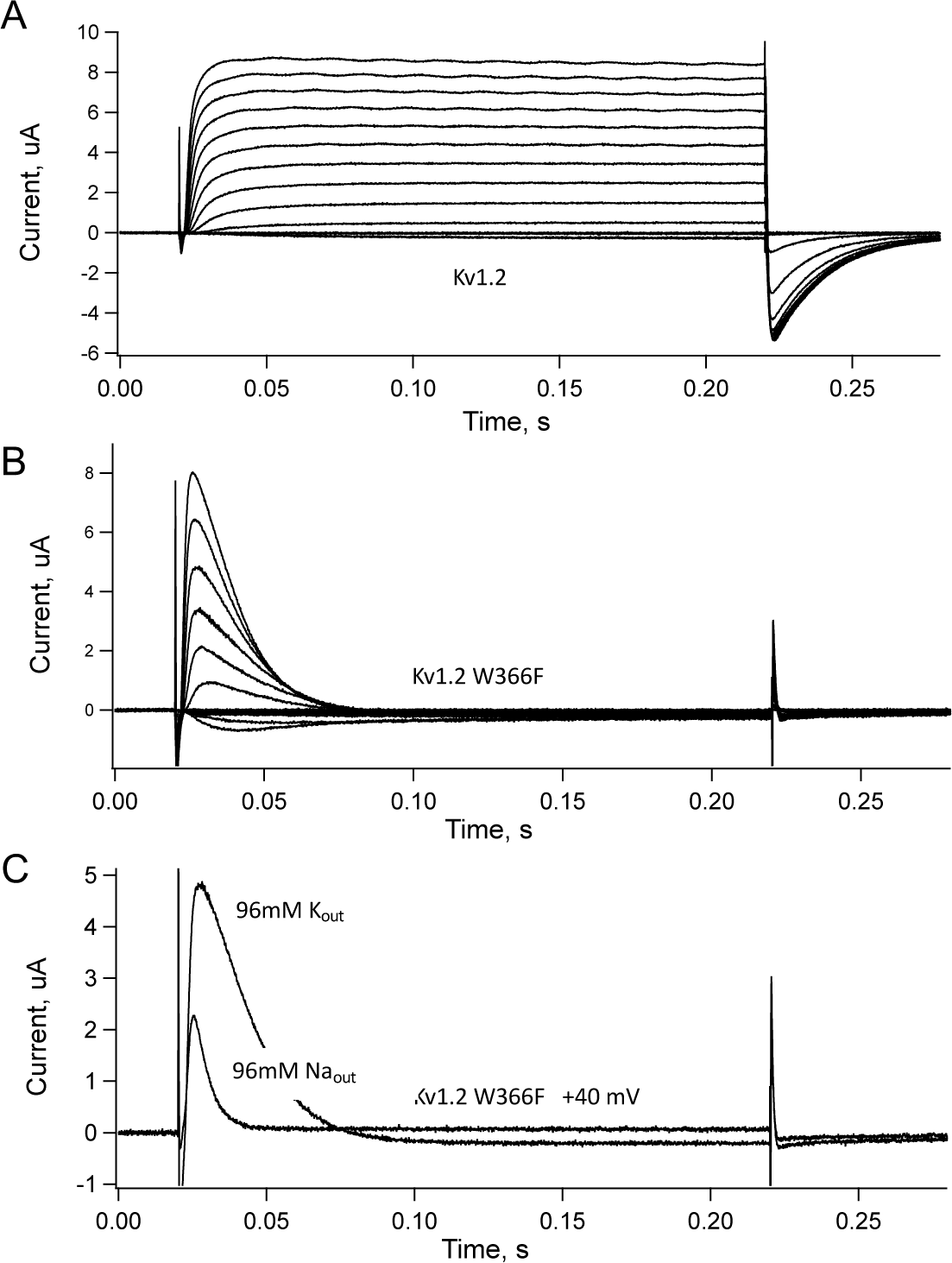
currents from native and W366F Kv1.2 channels. *Xenopus* oocytes were injected with mRNA for the alpha subunit constructs used in this study. **A**, native Kv1.2 currents elicited from pulses to -60 to + 60mV in 10 mV steps, from a holding potential of -80mV. **B**, Same voltage protocol applied to channels with W366F alpha subunits, recorded with 96 mM K^+^ bath solution. **C**, Comparison of currents elicited from an oocyte at +40 mV with 96mM K^+^ or Na^+^ bath solutions. The potassium-free Na^+^ solution yielded faster inactivation, as expected for C-type inactivation. The apparently sustained current in 95 mM Na^+^ solution is an artifact of P/4 leak subtraction at -120mV holding potential.

**Figure 2 - figure supplement 2.**
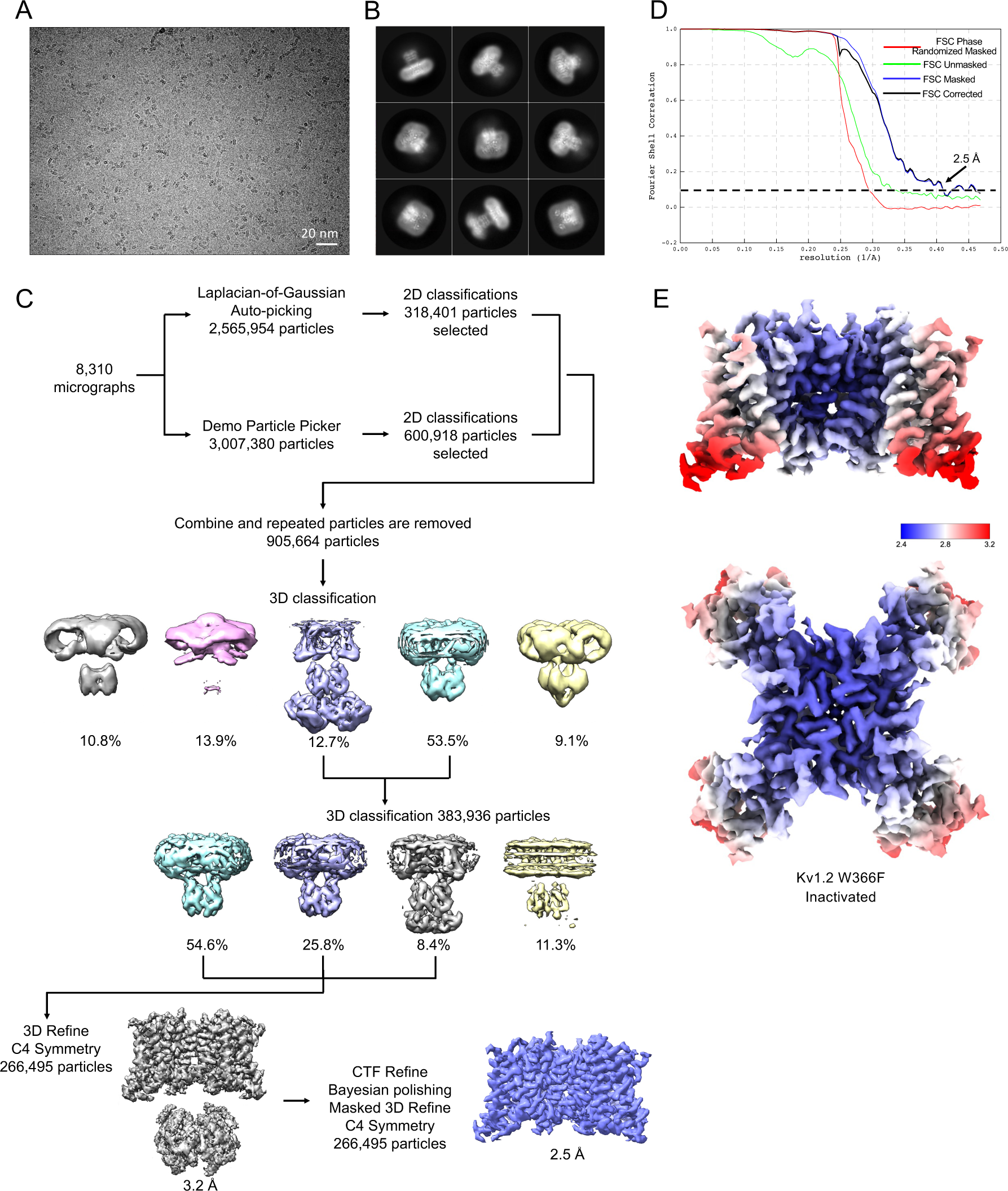
Processing of Kv1.2 W366F images. (A) Representative micrograph. (B) Representative 2D classes. (C) Cryo-EM data processing workflow. (D) Gold standard FSC resolution estimation. (F) Local resolution estimation.

**Figure 2 - figure supplement 3.**
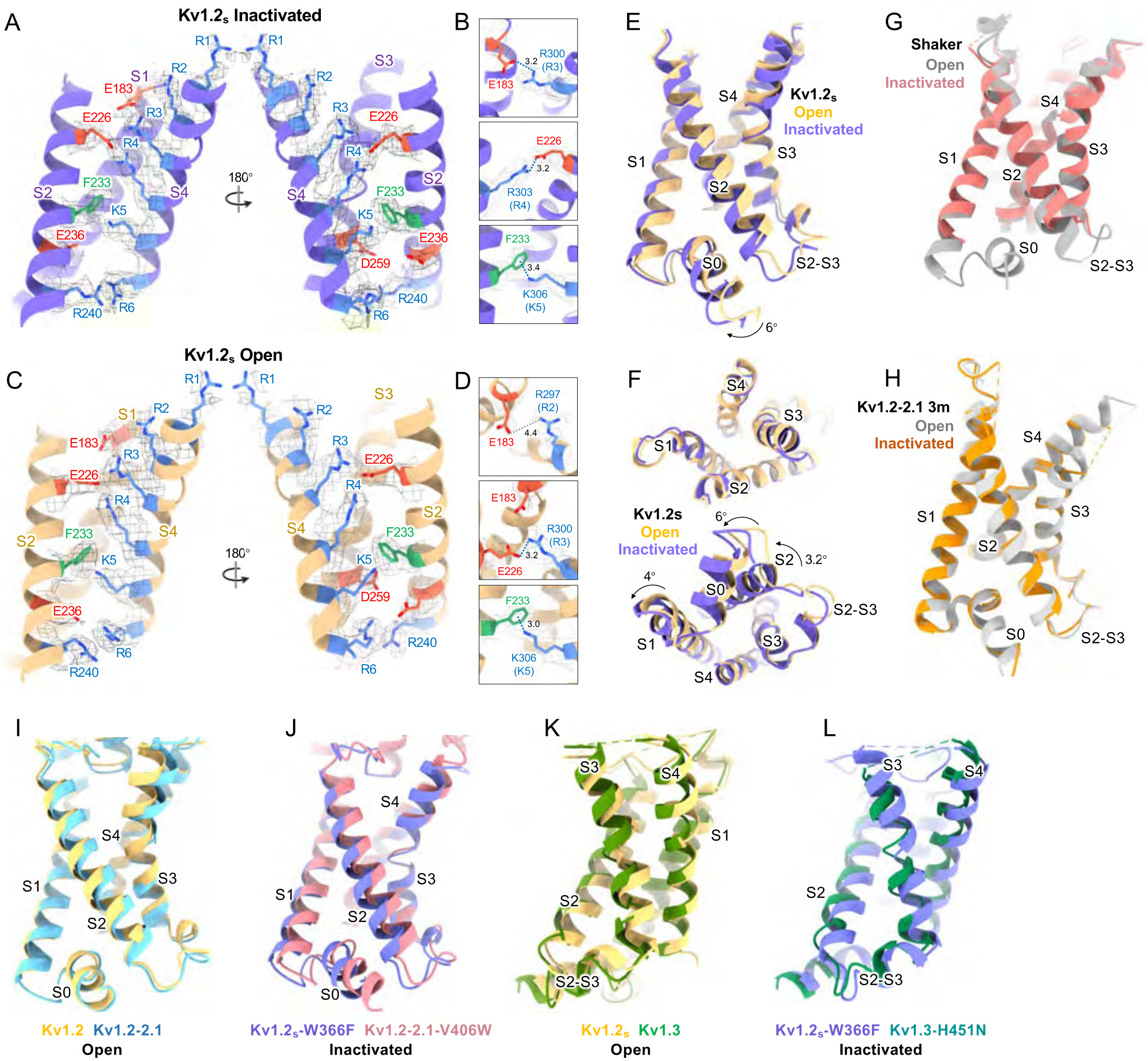
Voltage-sensing-domain conformational differences between open and C-type inactivated states. (A) Side view of VSD structure and maps of Kv1.2_s_ in the inactivated state. (B) Close-up of R3/E183 (upper), R4/E226 (middle), and K5/F233 (lower) interactions in the inactivated state. (C) Kv1.2_s_ VSD structure in open state. (D) Close-up of R2/E138 (upper) and R3/E226 (middle), K5/F233 (lower) interactions in the open state. Side view (E), top view (F, upper) and bottom view (F, lower) of the VSD conformational difference between open (yellow) and inactivated (purple) states. Superposition of (G) Shaker open (PDB: 7SIP), Shaker W434F inactivated (PDB: 7SJ1) and (F) Kv1.2-2.1open (PDB: 7SIZ), Kv1.2-2.1 3minactivated (PDB: 7SIT) VSD structures. (I-L) Superposition of VSD structures. (I) Kv1.2_s_ and Kv1.2-2.1 (PDB: 2R9R); (J) Kv1.2_s_ W366Fand Kv1.2-2.1 V406W (PDB: 5WIE); (K) Kv1.2and Kv1.3 (PDB: 7EJ1); (L) Kv1.2_s_ W366Fand Kv1.3 H451N (PDB: 7EJ2).

**Figure 2 - figure supplement 4.**
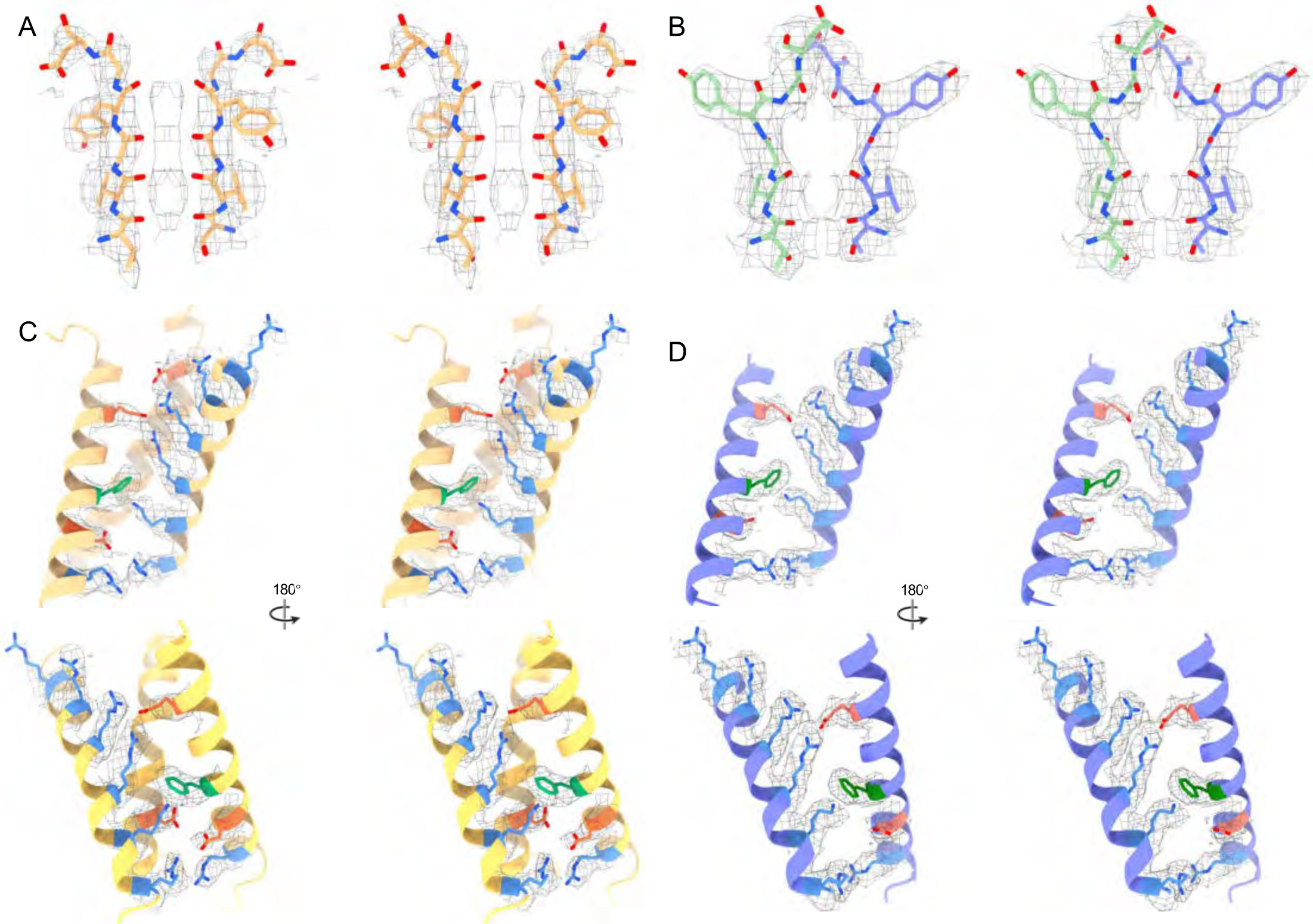
Stereo views of selectivity filter and voltage-sensing-domain. Stereo view of (A) Kv1.2s SF, (B) Kv1.2s-W366F SF, (C) Kv1.2s VSD, (D) Kv1.2s-W366F VSD.

**Figure 3 - figure supplement 1.**
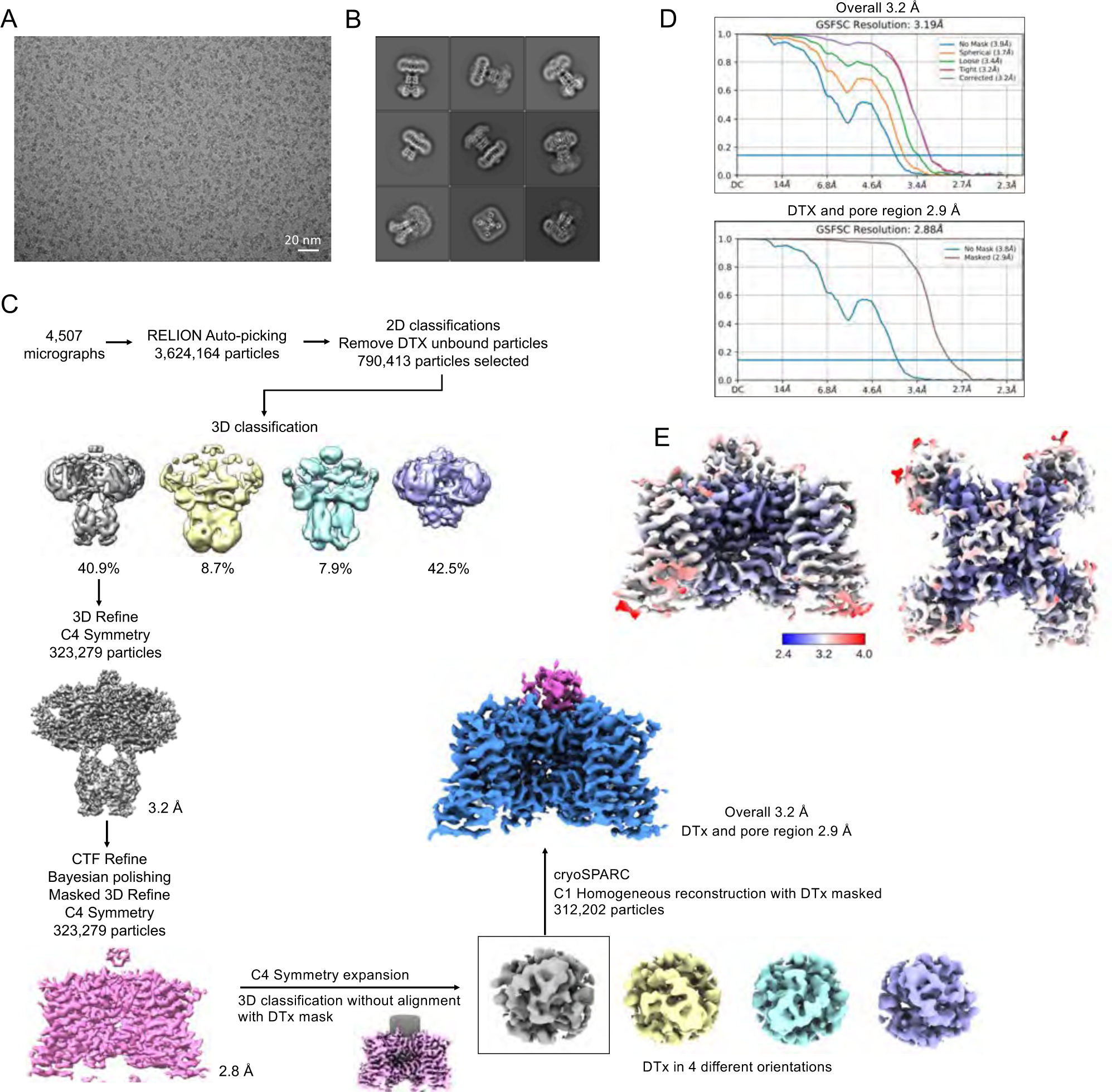
Cryo-EM imaging and reconstruction of Kv1.2_s_-DTX. (A) Representative micrograph. (B) Representative 2D classes, showing the DTx “cap” on the particles. (C) Cryo-EM data processing workflow. See Methods for details of the symmetry expansion and C1 reconstruction. (D) Gold standard FSC resolution estimation for the overall map (top) and the DTX-plus-selectivity filter masked region. (E) Local resolution estimation.

**Figure 3 - figure supplement 2.**
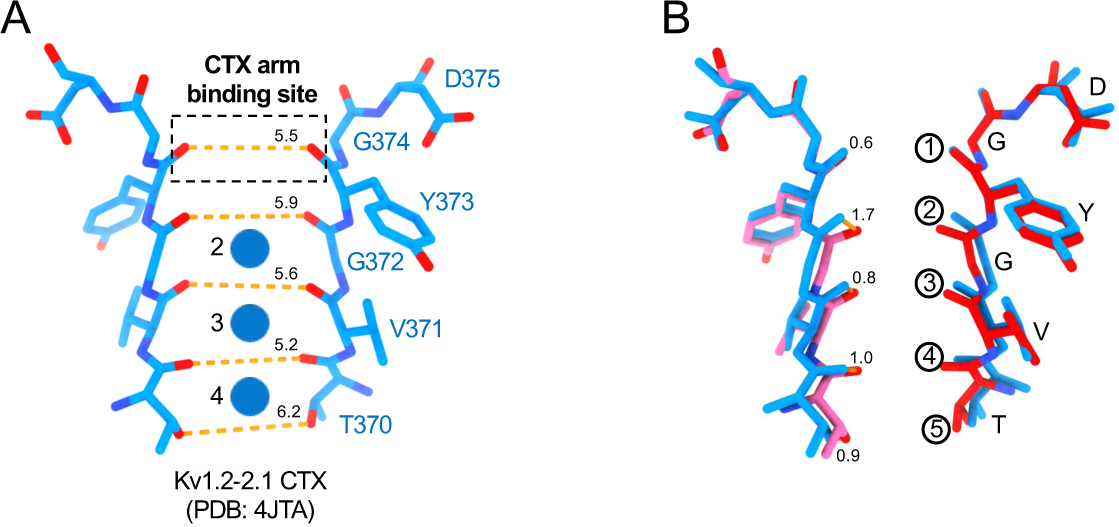
Comparison of the Kv1.2-2.1 CTx bound selectivity filter with the Kv1.2_s_ DTx-bound structure. (A) Side view of the selectivity filter of Kv1.2-2.1 CTx bound conformation (Banerjee et al. 2013). Orange dashed lines show the distances between carbonyl oxygens, for comparison with Fig. 3G. Potassium ions are shown as blue balls. (B) Superposition of Kv1.2 DTx (red) and Kv1.2-2.1 CTx (blue) selectivity filter structures. Apparent carbonyl displacements are given in Å.

**Figure 4 - figure supplement 1.**
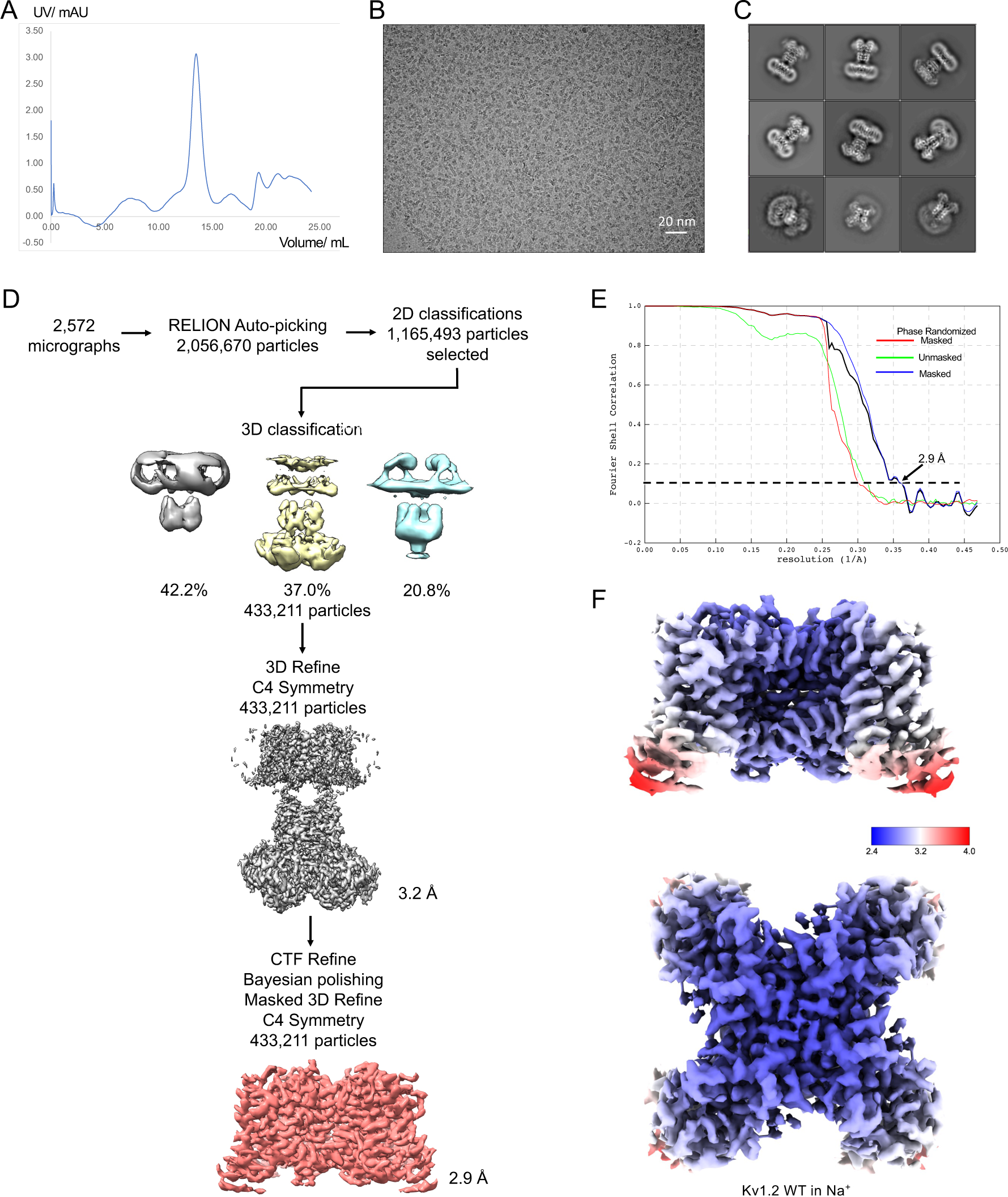
Cryo-EM of Kv1.2_s_ in Na^+^. (A) Size-exclusion chromatogram. Detector drift was large compared to a small protein signal. (B) Representative micrograph showing monodisperse particles on the graphene substrate. (C) Representative 2D classes. (D) Cryo-EM data processing workflow. (E) Gold standard FSC resolution estimation. (F) Local resolution estimation.

**Figure 4 - figure supplement 2.**
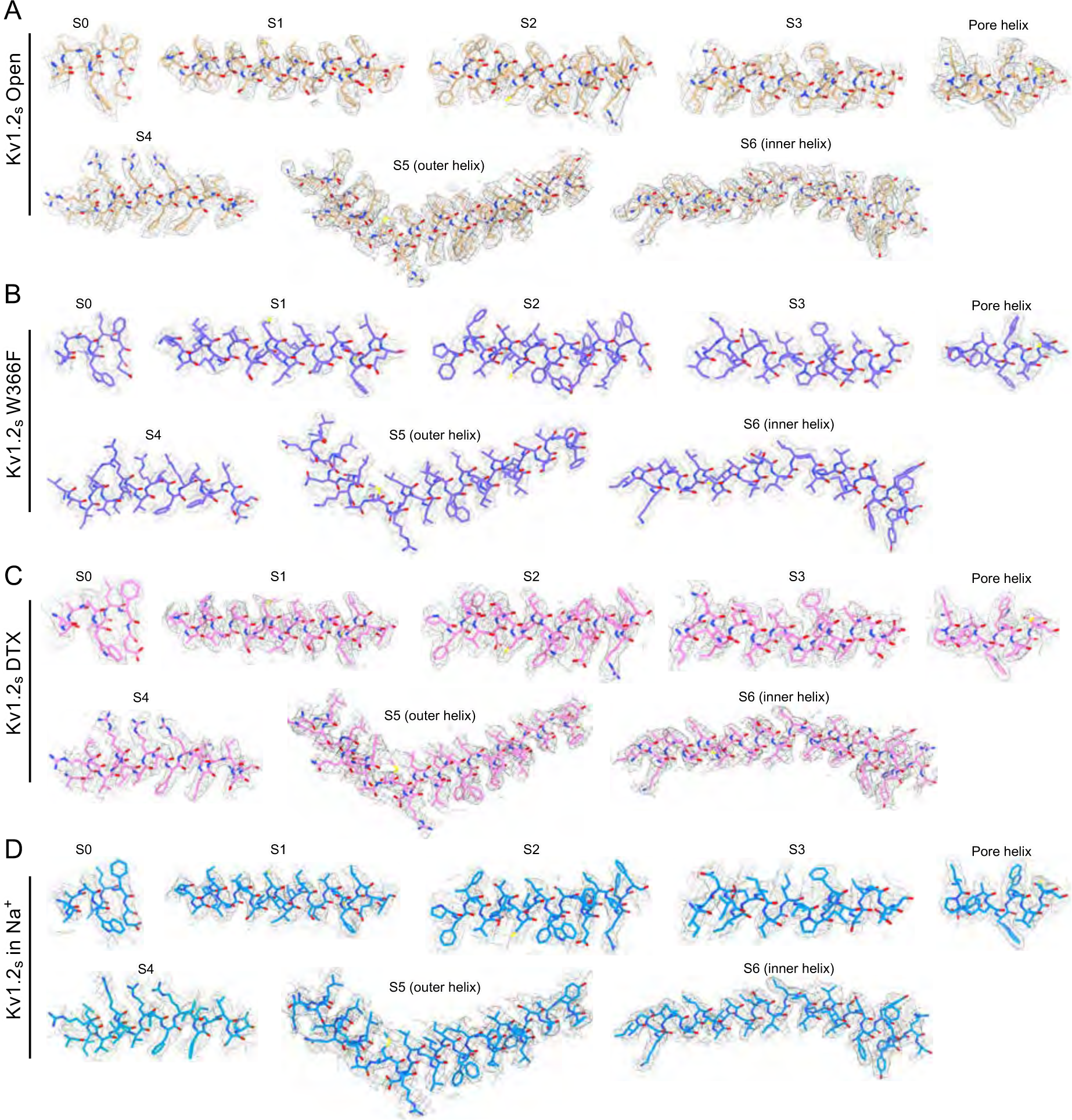
Comparisons of cryo-EM density map and model for alpha helices in each Kv1.2 structure reported here.

**Figure 4 - figure supplement 3.**
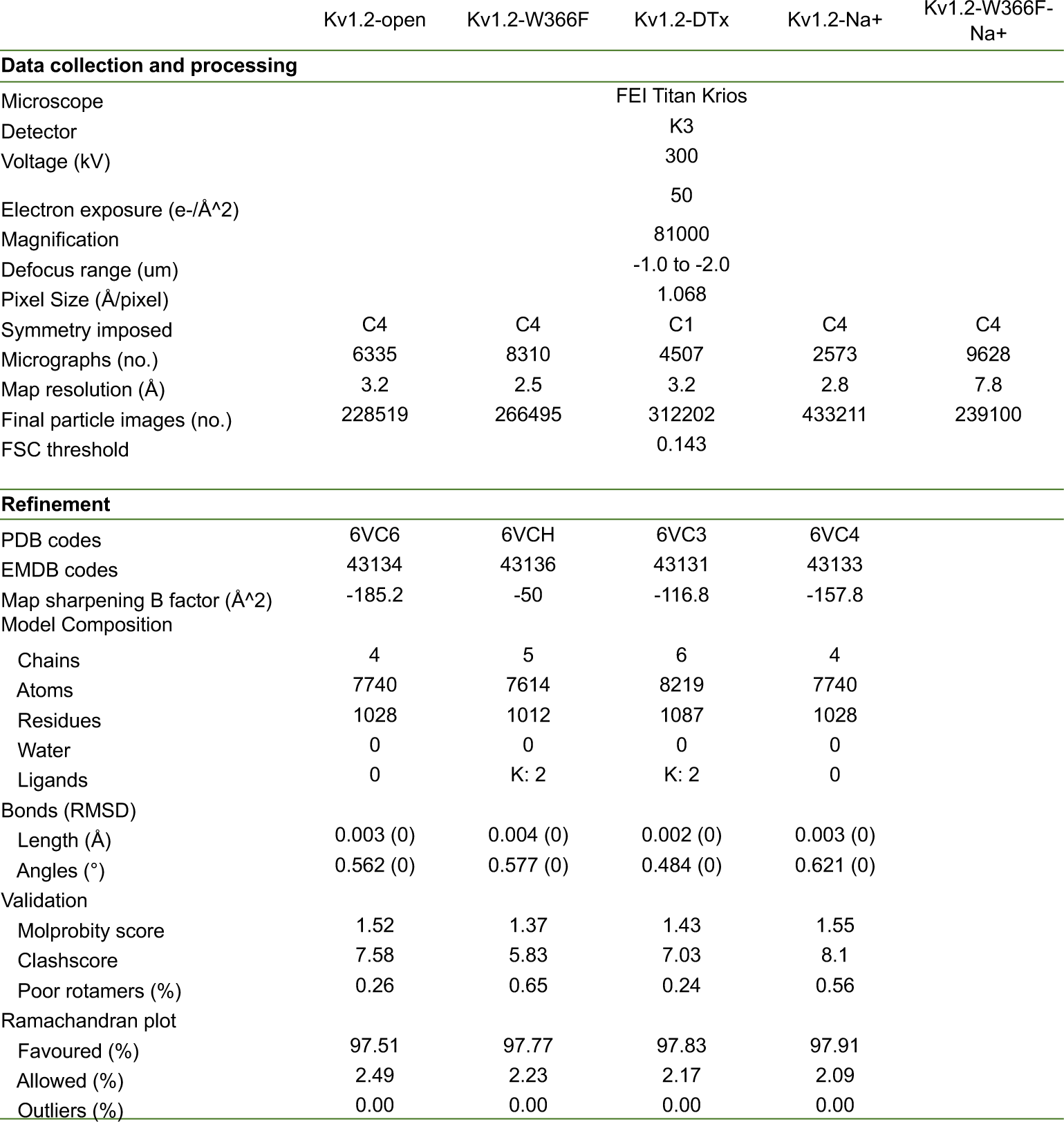
Cryo-EM data collection, refinement and validation statistics

**Figure 5 - figure supplement 1.**
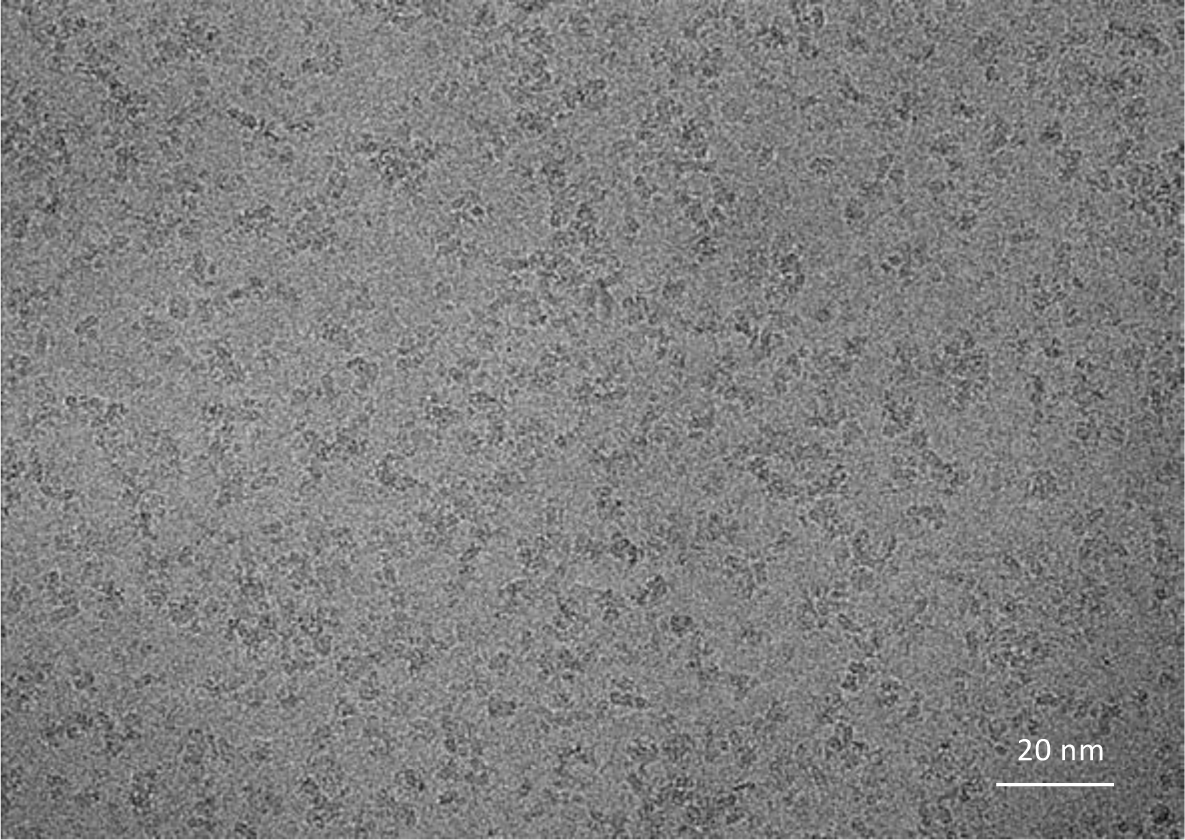
Representative micrograph of Kv1.2 W366F in Na^+^, demonstrating the absence of protein aggregates on the graphene substrate.

**Figure 6–Figure Supplement 1.**
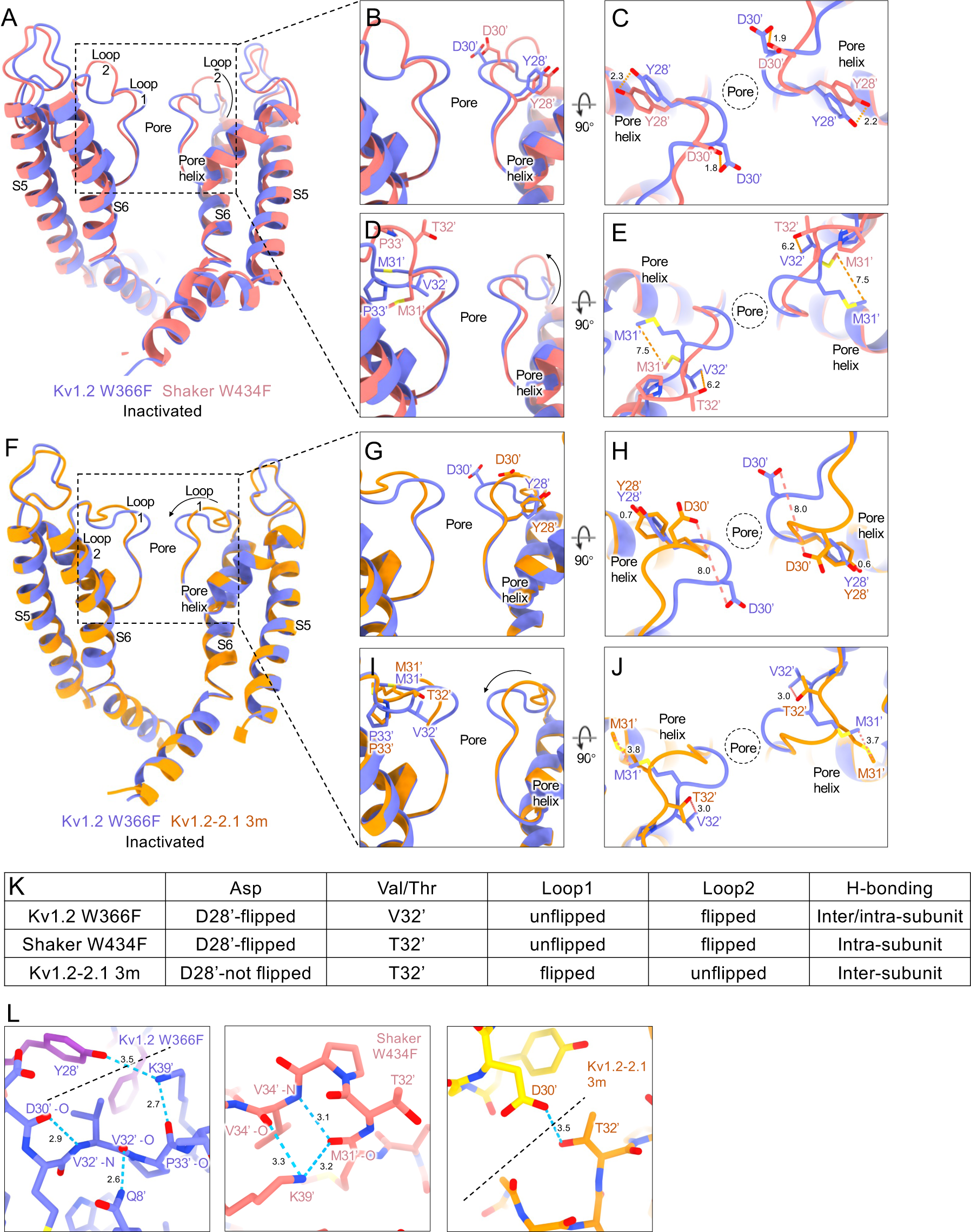
Structural comparison of inactivated Kv channels. (A-J) Structural superposition of Kv1.2_s_-W366F pore domain with other inactivated channels: side view with Shaker W434F (A) or Kv1.2-2.1-3m (F). Loop 1 conformational differences with Shaker W434F side view (B), top view (C); or with Kv1.2-2.1-3m side view (G), top view (H). Loop 2 conformational changes with Shaker W434F side view (D), top view (E); Kv1.2-2.1 3m side view (I), top view (J). (K) Table lists of the differences among the inactivated Kv channels. (L) H-bonds among the inactivated Kv channels. Adjacent subunits are shown as different colors, and a dashed black line denotes the subunit boundary. H-bonding patterns affect the stability of the inactivated state and an obvious difference is at the location 32’, where Shaker has Thr and Kv1.2 has Val. The mutation V32’T in the Kv1.2-2.1 background provides a new hydrogen bond that stabilizes the important residue D30’ in the inactivated conformation; this can be seen in the Kv1.2-2.1-3m structure (L, panel 3). Our inactivated Kv1.2s structure (containing V32’) nevertheless shows an H-bond network that includes Y28’ and D30’ along with main-chain atoms, possibly yielding a similar stabilization of the inactivated state (L, panel 1). The inactivated Shaker structure lacks H-bond partners for either Y28’ or D30’, which instead are exposed to solvent; however another H-bond network stabilizes the P-loop-S6 linker (L panel 2).

**Figure 6 - figure supplement 2.**
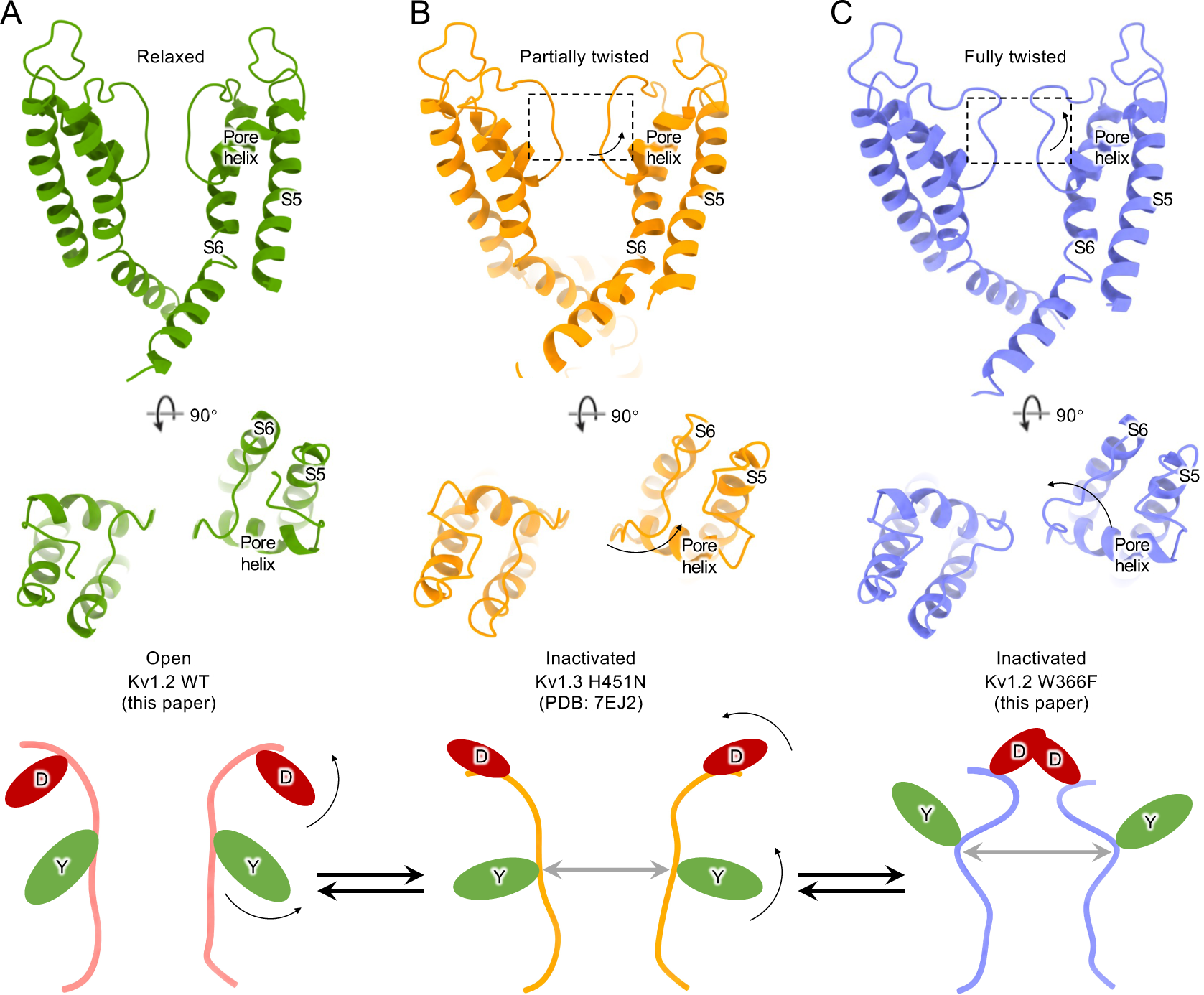
Summary of conformational changes in Kv channel inactivation. Upper panels: (A) Kv1.2WT pore domain (PD) in green (B) Kv1.3 H451N PD in orange and (C) Kv1.2 W366F PD in orchid represent the relaxed, partially twisted and fully twisted P-loop respectively. Lower panels: cartoon illustration of (A) relaxed, (B) partially twisted and (C) fully twisted selectivity filter P-loop of Kv channels. D30’ and Y28’ residue side chains are shown as red and green ovals.

## References

Banerjee, A., Lee, A., Campbell, E., & Mackinnon, R. (2013). Structure of a pore-blocking toxin in complex with a eukaryotic voltage-dependent K(+) channel. Elife, 2, e00594. 10.7554/eLife.00594

Bassetto, C. A., Carvalho-de-Souza, J. L., & Bezanilla, F. (2021). Molecular basis for functional connectivity between the voltage sensor and the selectivity filter gate in Shaker K(+) channels. Elife, 10. 10.7554/eLife.63077

Conti, L., Renhorn, J., Gabrielsson, A., Turesson, F., Liin, S. I., Lindahl, E., & Elinder, F. (2016). Reciprocal voltage sensor-to-pore coupling leads to potassium channel C-type inactivation. Sci Rep, 6, 27562. 10.1038/srep27562

Cuello, L. G., Cortes, D. M., & Perozo, E. (2017). The gating cycle of a K(+) channel at atomic resolution. Elife, 6. 10.7554/eLife.28032

Doyle, D. A., Morais Cabral, J., Pfuetzner, R. A., Kuo, A., Gulbis, J. M., Cohen, S. L., Chait, B. T., & MacKinnon, R. (1998). The structure of the potassium channel: molecular basis of K+ conduction and selectivity. Science, 280(5360), 69–77. 10.1126/science.280.5360.69

Emsley, P., Lohkamp, B., Scott, W. G., & Cowtan, K. (2010). Features and development of Coot. Acta Crystallogr D Biol Crystallogr, 66(Pt 4), 486–501. 10.1107/S0907444910007493

Fan, X., Wang, J., Zhang, X., Yang, Z., Zhang, J. C., Zhao, L., Peng, H. L., Lei, J., & Wang, H. W. (2019). Single particle cryo-EM reconstruction of 52 kDa streptavidin at 3.2 Angstrom resolution. Nat Commun, 10(1), 2386. 10.1038/s41467-019-10368-w

Gasparini, S., Danse, J. M., Lecoq, A., Pinkasfeld, S., Zinn-Justin, S., Young, L. C., de Medeiros, C. C., Rowan, E. G., Harvey, A. L., & Menez, A. (1998). Delineation of the functional site of alpha-dendrotoxin. The functional topographies of dendrotoxins are different but share a conserved core with those of other Kv1 potassium channel-blocking toxins. J Biol Chem, 273(39), 25393-25403. 10.1074/jbc.273.39.25393

Goddard, T. D., Huang, C. C., Meng, E. C., Pettersen, E. F., Couch, G. S., Morris, J. H., & Ferrin, T. E. (2018). UCSF ChimeraX: Meeting modern challenges in visualization and analysis. Protein Sci, 27(1), 14–25. 10.1002/pro.3235

Gulbis, J. M., Mann, S., & MacKinnon, R. (1999). Structure of a voltage-dependent K+ channel beta subunit. Cell, 97(7), 943–952. 10.1016/s0092-8674(00)80805-3

Gulbis, J. M., Zhou, M., Mann, S., & MacKinnon, R. (2000). Structure of the cytoplasmic beta subunit-T1 assembly of voltage-dependent K+ channels. Science, 289(5476), 123–127. 10.1126/science.289.5476.123

Harvey, A. L., Rowan, E. G., Vatanpour, H., Engstrom, A., Westerlund, B., & Karlsson, E. (1997). Changes to biological activity following acetylation of dendrotoxin I from Dendroaspis polylepis (black mamba). Toxicon, 35(8), 1263–1273. 10.1016/s0041-0101(97)00016-0

Heginbotham, L., Lu, Z., Abramson, T., & MacKinnon, R. (1994). Mutations in the K+ channel signature sequence. Biophys J, 66(4), 1061–1067. 10.1016/S0006-3495(94)80887-2

Hodgkin, A. L., & Huxley, A. F. (1952). The components of membrane conductance in the giant axon of Loligo. J Physiol, 116(4), 473–496. 10.1113/jphysiol.1952.sp004718

Hoshi, T., Zagotta, W. N., & Aldrich, R. W. (1991). Two types of inactivation in Shaker K+ channels: effects of alterations in the carboxy-terminal region. Neuron, 7(4), 547–556. 10.1016/0896-6273(91)90367-9

Ishida, I. G., Rangel-Yescas, G. E., Carrasco-Zanini, J., & Islas, L. D. (2015). Voltage-dependent gating and gating charge measurements in the Kv1.2 potassium channel. J Gen Physiol, 145(4), 345–358. 10.1085/jgp.201411300

Islas, L. D. (2016). Functional diversity of potassium channel voltage-sensing domains. Channels (Austin), 10(3), 202–213. 10.1080/19336950.2016.1141842

Iverson, L. E., Tanouye, M. A., Lester, H. A., Davidson, N., & Rudy, B. (1988). A-type potassium channels expressed from Shaker locus cDNA. Proc Natl Acad Sci U S A, 85(15), 5723–5727. 10.1073/pnas.85.15.5723

Jensen, M. O., Jogini, V., Borhani, D. W., Leffler, A. E., Dror, R. O., & Shaw, D. E. (2012). Mechanism of voltage gating in potassium channels. Science, 336(6078), 229–233. 10.1126/science.1216533

Karbat, I., Altman-Gueta, H., Fine, S., Szanto, T., Hamer-Rogotner, S., Dym, O., Frolow, F., Gordon, D., Panyi, G., Gurevitz, M., & Reuveny, E. (2019). Pore-modulating toxins exploit inherent slow inactivation to block K(+) channels. Proc Natl Acad Sci U S A, 116(37), 18700–18709. 10.1073/pnas.1908903116

Kondo, H. X., Yoshida, N., Shirota, M., & Kinoshita, K. (2018). Molecular Mechanism of Depolarization-Dependent Inactivation in W366F Mutant of Kv1.2. J Phys Chem B, 122(48), 10825–10833. 10.1021/acs.jpcb.8b09446

Korn, S. J., & Ikeda, S. R. (1995). Permeation selectivity by competition in a delayed rectifier potassium channel. Science, 269(5222), 410–412. 10.1126/science.7618108

Lee, C. H., & MacKinnon, R. (2017). Structures of the Human HCN1 Hyperpolarization-Activated Channel. Cell, 168(1-2), 111–120 e111. 10.1016/j.cell.2016.12.023

Lee, S. Y., & MacKinnon, R. (2004). A membrane-access mechanism of ion channel inhibition by voltage sensor toxins from spider venom. Nature, 430(6996), 232–235. 10.1038/nature02632

Levy, D. I., & Deutsch, C. (1996). Recovery from C-type inactivation is modulated by extracellular potassium. Biophys J, 70(2), 798–805. 10.1016/S0006-3495(96)79619-4

Li, J., Ostmeyer, J., Cuello, L. G., Perozo, E., & Roux, B. (2018). Rapid constriction of the selectivity filter underlies C-type inactivation in the KcsA potassium channel. J Gen Physiol, 150(10), 1408–1420. 10.1085/jgp.201812082

Li, J., Shen, R., Reddy, B., Perozo, E., & Roux, B. (2021). Mechanism of C-type inactivation in the hERG potassium channel. Sci Adv, 7(5). 10.1126/sciadv.abd6203

Li, J., Shen, R., Rohaim, A., Mendoza Uriarte, R., Fajer, M., Perozo, E., & Roux, B. (2021). Computational study of non-conductive selectivity filter conformations and C-type inactivation in a voltage-dependent potassium channel. J Gen Physiol, 153(9). 10.1085/jgp.202112875

Liebschner, D., Afonine, P. V., Baker, M. L., Bunkoczi, G., Chen, V. B., Croll, T. I., Hintze, B., Hung, L. W., Jain, S., McCoy, A. J., Moriarty, N. W., Oeffner, R. D., Poon, B. K., Prisant, M. G., Read, R. J., Richardson, J. S., Richardson, D. C., Sammito, M. D., Sobolev, O. V., . . . Adams, P. D. (2019). Macromolecular structure determination using X-rays, neutrons and electrons: recent developments in Phenix. Acta Crystallogr D Struct Biol, 75(Pt 10), 861–877. 10.1107/S2059798319011471

Liu, S., Zhao, Y., Dong, H., Xiao, L., Zhang, Y., Yang, Y., Ong, S. T., Chandy, K. G., Zhang, L., & Tian, C. (2021). Structures of wild-type and H451N mutant human lymphocyte potassium channel KV1.3. Cell Discov, 7(1), 39. 10.1038/s41421-021-00269-y

Lolicato, M., Natale, A. M., Abderemane-Ali, F., Crottes, D., Capponi, S., Duman, R., Wagner, A., Rosenberg, J. M., Grabe, M., & Minor, D. L., Jr. (2020). K(2P) channel C-type gating involves asymmetric selectivity filter order-disorder transitions. Sci Adv, 6(44). 10.1126/sciadv.abc9174

Long, S. B., Campbell, E. B., & Mackinnon, R. (2005). Crystal structure of a mammalian voltage-dependent Shaker family K+ channel. Science, 309(5736), 897–903. 10.1126/science.1116269

Long, S. B., Tao, X., Campbell, E. B., & MacKinnon, R. (2007). Atomic structure of a voltage-dependent K+ channel in a lipid membrane-like environment. Nature, 450(7168), 376–382. 10.1038/nature06265

Matamoros, M., Ng, X. W., Brettmann, J. B., Piston, D. W., & Nichols, C. G. (2023). Conformational plasticity of NaK2K and TREK2 potassium channel selectivity filters. Nat Commun, 14(1), 89. 10.1038/s41467-022-35756-7

Matthies, D., Bae, C., Toombes, G. E., Fox, T., Bartesaghi, A., Subramaniam, S., & Swartz, K. J. (2018). Single-particle cryo-EM structure of a voltage-activated potassium channel in lipid nanodiscs. Elife, 7. 10.7554/eLife.37558

Melishchuk, A., Loboda, A., & Armstrong, C. M. (1998). Loss of shaker K channel conductance in 0 K+ solutions: role of the voltage sensor. Biophys J, 75(4), 1828–1835. 10.1016/S0006-3495(98)77624-6

Mita, K., Sumikama, T., Iwamoto, M., Matsuki, Y., Shigemi, K., & Oiki, S. (2021). Conductance selectivity of Na(+) across the K(+) channel via Na(+) trapped in a tortuous trajectory. Proc Natl Acad Sci U S A, 118(12). 10.1073/pnas.2017168118

Morais-Cabral, J. H., Zhou, Y., & MacKinnon, R. (2001). Energetic optimization of ion conduction rate by the K+ selectivity filter. Nature, 414(6859), 37–42. 10.1038/35102000

Murshudov, G. N., Skubak, P., Lebedev, A. A., Pannu, N. S., Steiner, R. A., Nicholls, R. A., Winn, M. D., Long, F., & Vagin, A. A. (2011). REFMAC5 for the refinement of macromolecular crystal structures. Acta Crystallogr D Biol Crystallogr, 67(Pt 4), 355–367. 10.1107/S0907444911001314

Noskov, S. Y., & Roux, B. (2006). Ion selectivity in potassium channels. Biophys Chem, 124(3), 279–291. 10.1016/j.bpc.2006.05.033

Papazian, D. M., Schwarz, T. L., Tempel, B. L., Jan, Y. N., & Jan, L. Y. (1987). Cloning of genomic and complementary DNA from Shaker, a putative potassium channel gene from Drosophila. Science, 237(4816), 749–753. 10.1126/science.2441470

Perozo, E., MacKinnon, R., Bezanilla, F., & Stefani, E. (1993). Gating currents from a nonconducting mutant reveal open-closed conformations in Shaker K+ channels. Neuron, 11(2), 353–358. 10.1016/0896-6273(93)90190-3

Pettersen, E. F., Goddard, T. D., Huang, C. C., Couch, G. S., Greenblatt, D. M., Meng, E. C., & Ferrin, T. E. (2004). UCSF Chimera--a visualization system for exploratory research and analysis. J Comput Chem, 25(13), 1605–1612. 10.1002/jcc.20084

Pless, S. A., Galpin, J. D., Niciforovic, A. P., Kurata, H. T., & Ahern, C. A. (2013). Hydrogen bonds as molecular timers for slow inactivation in voltage-gated potassium channels. Elife, 2, e01289. 10.7554/eLife.01289

Reddi, R., Matulef, K., Riederer, E. A., Whorton, M. R., & Valiyaveetil, F. I. (2022). Structural basis for C-type inactivation in a Shaker family voltage-gated K(+) channel. Sci Adv, 8(16), eabm8804. 10.1126/sciadv.abm8804

Rettig, J., Heinemann, S. H., Wunder, F., Lorra, C., Parcej, D. N., Dolly, J. O., & Pongs, O. (1994). Inactivation properties of voltage-gated K+ channels altered by presence of beta-subunit. Nature, 369(6478), 289–294. 10.1038/369289a0

Rosenthal, J. J., & Bezanilla, F. (2002). A comparison of propagated action potentials from tropical and temperate squid axons: different durations and conduction velocities correlate with ionic conductance levels. J Exp Biol, 205(Pt 12), 1819–1830. 10.1242/jeb.205.12.1819

Roux, B. (2005). Ion conduction and selectivity in K(+) channels. Annu Rev Biophys Biomol Struct, 34, 153–171. 10.1146/annurev.biophys.34.040204.144655

Sauer, D. B., Zeng, W., Raghunathan, S., & Jiang, Y. (2011). Protein interactions central to stabilizing the K+ channel selectivity filter in a four-sited configuration for selective K+ permeation. Proc Natl Acad Sci U S A, 108(40), 16634–16639. 10.1073/pnas.1111688108

Schorb, M., Haberbosch, I., Hagen, W. J. H., Schwab, Y., & Mastronarde, D. N. (2019). Software tools for automated transmission electron microscopy. Nat Methods, 16(6), 471–477. 10.1038/s41592-019-0396-9

Selvakumar, P., Fernandez-Marino, A. I., Khanra, N., He, C., Paquette, A. J., Wang, B., Huang, R., Smider, V. V., Rice, W. J., Swartz, K. J., & Meyerson, J. R. (2022). Structures of the T cell potassium channel Kv1.3 with immunoglobulin modulators. Nat Commun, 13(1), 3854. 10.1038/s41467-022-31285-5

Shealy, R. T., Murphy, A. D., Ramarathnam, R., Jakobsson, E., & Subramaniam, S. (2003). Sequence-function analysis of the K+-selective family of ion channels using a comprehensive alignment and the KcsA channel structure. Biophys J, 84(5), 2929–2942. 10.1016/S0006-3495(03)70020-4

Shi, N., Ye, S., Alam, A., Chen, L., & Jiang, Y. (2006). Atomic structure of a Na+- and K+-conducting channel. Nature, 440(7083), 570–574. 10.1038/nature04508

Skarzynski, T. (1992). Crystal structure of alpha-dendrotoxin from the green mamba venom and its comparison with the structure of bovine pancreatic trypsin inhibitor. J Mol Biol, 224(3), 671–683. 10.1016/0022-2836(92)90552-u

Starkus, J. G., Kuschel, L., Rayner, M. D., & Heinemann, S. H. (1997). Ion conduction through C-type inactivated Shaker channels. J Gen Physiol, 110(5), 539–550. 10.1085/jgp.110.5.539

Starkus, J. G., Kuschel, L., Rayner, M. D., & Heinemann, S. H. (1998). Macroscopic Na+ currents in the “Nonconducting” Shaker potassium channel mutant W434F. J Gen Physiol, 112(1), 85–93. 10.1085/jgp.112.1.85

Suarez-Delgado, E., Rangel-Sandin, T. G., Ishida, I. G., Rangel-Yescas, G. E., Rosenbaum, T., & Islas, L. D. (2020). KV1.2 channels inactivate through a mechanism similar to C-type inactivation. J Gen Physiol, 152(6). 10.1085/jgp.201912499

Tan, X. F., Bae, C., Stix, R., Fernandez-Marino, A. I., Huffer, K., Chang, T. H., Jiang, J., Faraldo-Gomez, J. D., & Swartz, K. J. (2022). Structure of the Shaker Kv channel and mechanism of slow C-type inactivation. Sci Adv, 8(11), eabm7814. 10.1126/sciadv.abm7814

Tao, X., Lee, A., Limapichat, W., Dougherty, D. A., & MacKinnon, R. (2010). A gating charge transfer center in voltage sensors. Science, 328(5974), 67–73. 10.1126/science.1185954

Tao, X., & MacKinnon, R. (2008). Functional analysis of Kv1.2 and paddle chimera Kv channels in planar lipid bilayers. J Mol Biol, 382(1), 24–33. 10.1016/j.jmb.2008.06.085

Tyagi, A., Ahmed, T., Jian, S., Bajaj, S., Ong, S. T., Goay, S. S. M., Zhao, Y., Vorobyov, I., Tian, C., Chandy, K. G., & Bhushan, S. (2022). Rearrangement of a unique Kv1.3 selectivity filter conformation upon binding of a drug. Proc Natl Acad Sci U S A, 119(5). 10.1073/pnas.2113536119

Wang, J. M., Roh, S. H., Kim, S., Lee, C. W., Kim, J. I., & Swartz, K. J. (2004). Molecular surface of tarantula toxins interacting with voltage sensors in K(v) channels. J Gen Physiol, 123(4), 455–467. 10.1085/jgp.200309005

Wang, S., Lee, S. J., Maksaev, G., Fang, X., Zuo, C., & Nichols, C. G. (2019). Potassium channel selectivity filter dynamics revealed by single-molecule FRET. Nat Chem Biol, 15(4), 377–383. 10.1038/s41589-019-0240-7

Wu, X., Gupta, K., & Swartz, K. J. (2022). Mutations within the selectivity filter reveal that Kv1 channels have distinct propensities to slow inactivate. J Gen Physiol, 154(11). 10.1085/jgp.202213222

Yang, Y., Yan, Y., & Sigworth, F. J. (1997). How does the W434F mutation block current in Shaker potassium channels? J Gen Physiol, 109(6), 779–789. 10.1085/jgp.109.6.779

Zhang, K. (2016). Gctf: Real-time CTF determination and correction. J Struct Biol, 193(1), 1–12. 10.1016/j.jsb.2015.11.003

Zhang, K., Julius, D., & Cheng, Y. (2021). Structural snapshots of TRPV1 reveal mechanism of polymodal functionality. Cell, 184(20), 5138–5150 e5112. 10.1016/j.cell.2021.08.012

Zheng, S. Q., Palovcak, E., Armache, J. P., Verba, K. A., Cheng, Y., & Agard, D. A. (2017). MotionCor2: anisotropic correction of beam-induced motion for improved cryo-electron microscopy. Nat Methods, 14(4), 331–332. 10.1038/nmeth.4193

Zhou, Y., Morais-Cabral, J. H., Kaufman, A., & MacKinnon, R. (2001). Chemistry of ion coordination and hydration revealed by a K+ channel-Fab complex at 2.0 A resolution. Nature, 414(6859), 43–48. 10.1038/35102009

